# A hybrid structure determination approach to investigate the druggability of the nucleocapsid protein of SARS-CoV-2

**DOI:** 10.1101/2022.09.15.507991

**Authors:** Giacomo Padroni, Maria Bikaki, Mihajlo Novakovic, Antje C. Wolter, Simon H. Rüdisser, Alvar D. Gossert, Alexander Leitner, Frederic H.-T Allain

**Affiliations:** Institute of Biochemistry, Department of Biology, ETH Zurich, Hönggerbergring 64, 8093 Zürich, Switzerland; Institute of Molecular Systems Biology, Department of Biology, ETH Zurich, Otto-Stern-Weg 3, 8093 Zürich, Switzerland; Biomolecular NMR Spectroscopy Platform, ETH Zurich, Hönggerbergring 64, 8093 Zurich, Switzerland

## Abstract

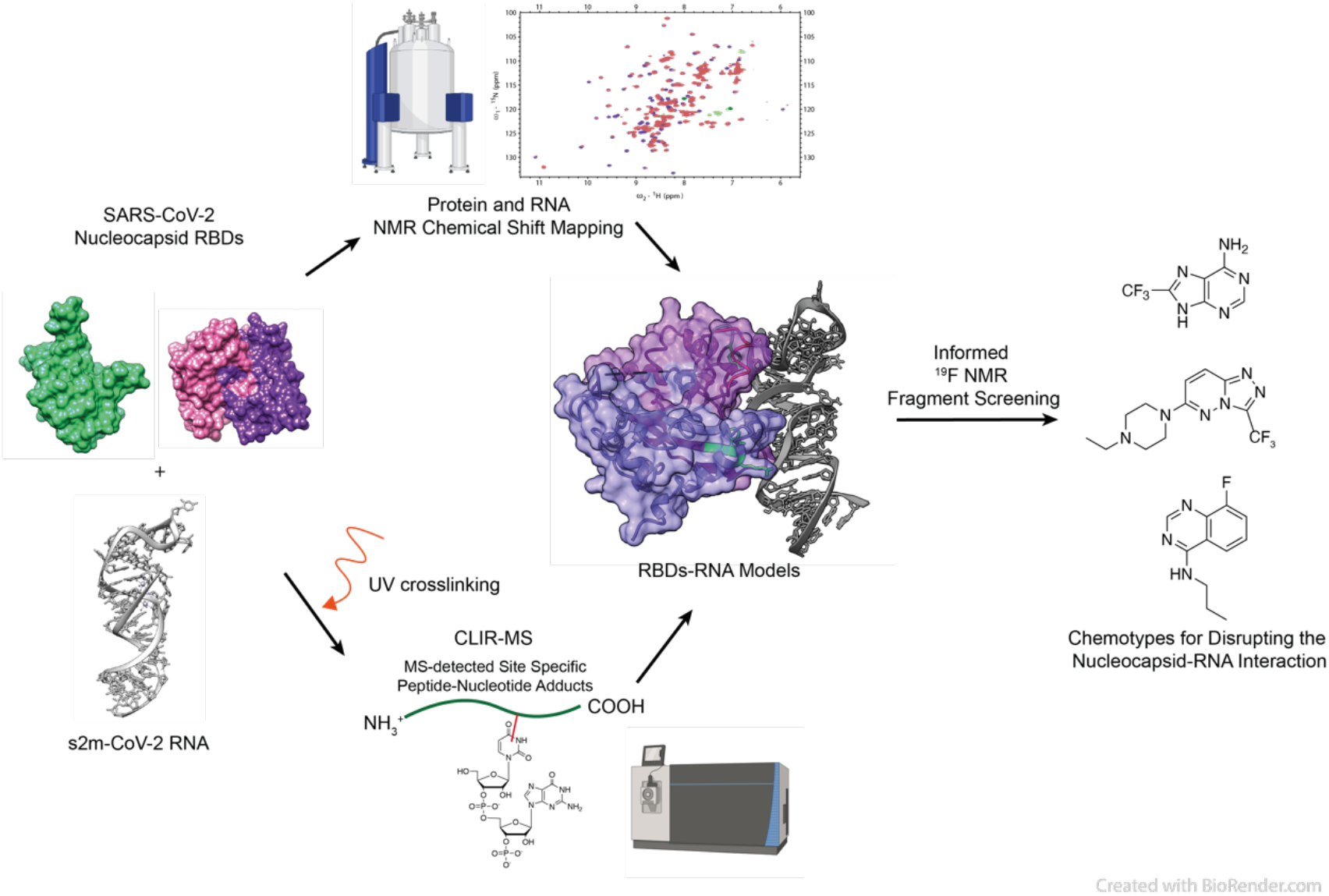

The ongoing pandemic caused by SARS-CoV-2 has called for concerted efforts to generate new insights into the biology of betacoronaviruses to inform drug screening and development. Here, we establish a workflow to determine the RNA recognition and druggability of the nucleocapsid N-protein of SARS-CoV-2, a highly abundant protein crucial for the viral life cycle. We use a synergistic method that combines NMR spectroscopy and protein-RNA cross-linking coupled to mass spectrometry to quickly determine the RNA binding of two RNA recognition domains of the N-protein. Finally, we explore the druggability of these domains by performing an NMR fragment screening. This workflow identified small molecule chemotypes that bind to RNA binding interfaces and that have promising properties for further drug development.

## INTRODUCTION

The coronavirus SARS-CoV-2 is an enveloped positive-sense single-stranded (ss) RNA virus responsible for the COVID-19 disease^1,2^. The viral RNA is approximately 30 kilobases long and is replicated and packaged in the host cells, a process that requires interactions with the viral RNA-binding proteins (RBP) and a number of host proteins^3,4^. While vaccine development has widely mitigated the severity of the disease, the investigation of such interactions is fundamental to understand disease progression and develop therapeutics to counteract viral variants that escape vaccine immunity. A crucial protein that mediates RNA packaging and replication is the nucleocapsid N-protein, a 45 kDa protein composed of two RNA binding domains (RBDs) flanked by intrinsically disordered regions (IDRs), including a serine/arginine-rich linker^5,6^ (Figure 1A). Currently, it is unclear how N-RNA interactions facilitate viral replication. The N-terminal RBD (NTD) interacts with RNA as a monomer and its interactions with ss and double-stranded (ds) RNA have been modelled^7^. Recent studies support the idea that despite being a promiscuous RNA binder, NTD binds to a variety of RNA structures with different signatures, suggesting a possible mechanism for regulation and RNA processing^8,9^. On the other hand, the C-terminal RBD (CTD) is present in solution as a swapped dimer as determined by x-ray crystallography, however little is known about the structural details of RNA recognition^10,11^. IDRs in the full-length protein are also known to promote liquid-liquid phase separation (LLPS) in the presence of RNA indicating an intricate system of protein-RNA interactions at the basis of the function of the N-protein^12–15^. Both CTD and NTD subdomains can also phase separate from a solution under a variety of external conditions^15,16^. Although the structural details of N-RNA recognition could be of crucial importance for the rational design of novel drugs, current techniques are limited by this Nucleocapsid condensation behaviour in the presence of RNA in solution, which precludes high-resolution structural determination.

**Figure 1.**
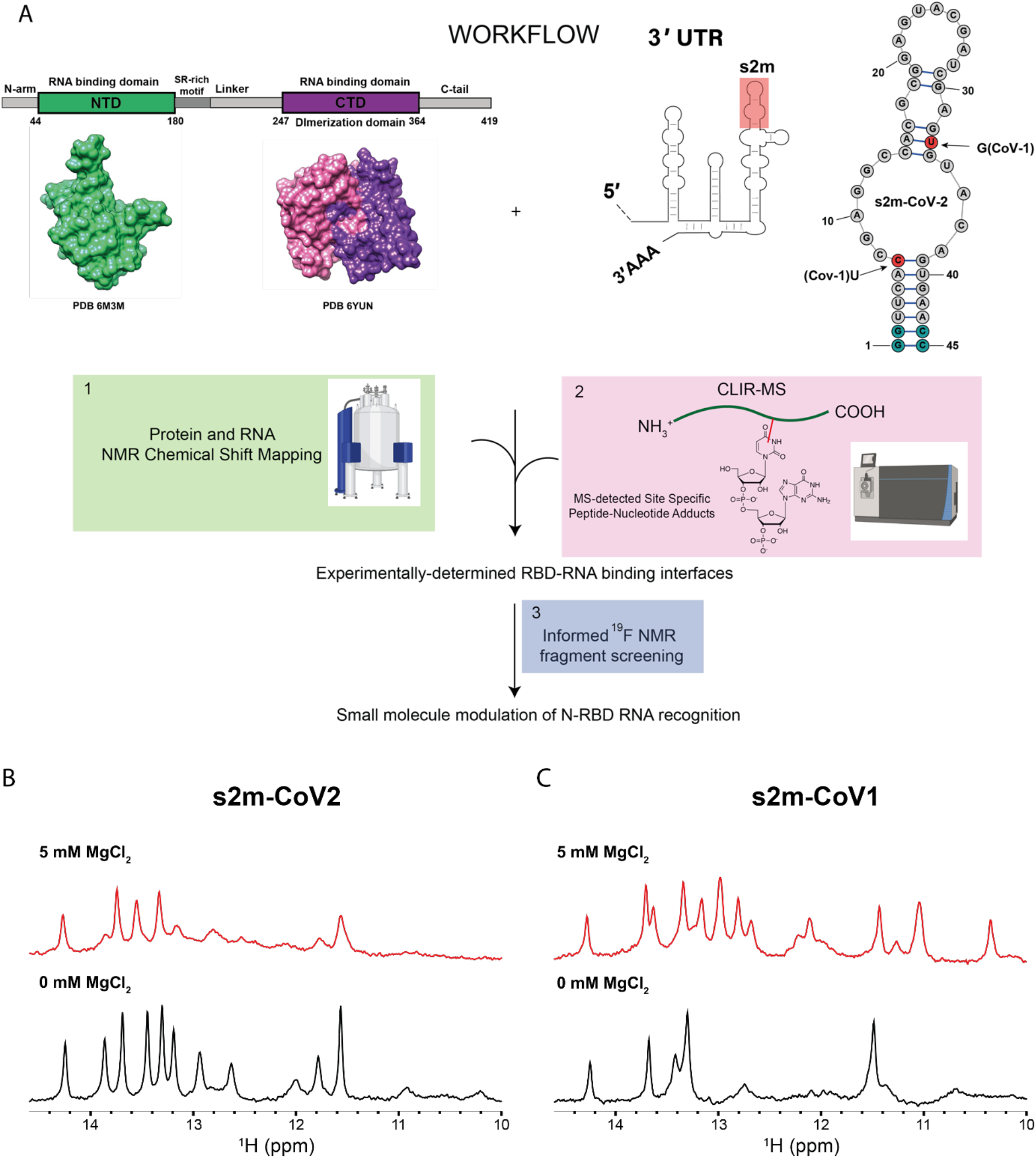
(A) Workflow including the schematic representation of N-protein primary structure and 3D structures of the RBDs (PDB 63M3 and 6YUN); the schematic representation of the 3’UTR region of SARS-CoV-2 genome highlighting the conserved RNA element s2m plus the schematic representation of the experimentally-determined secondary structure of s2m-CoV-2^20^ (red circles indicate nucleotide mutations between s2m-CoV-2 and s2m-CoV-1, light-blue circles indicate bases that were added to the construct to facilitate in vitro transcription). Created with BioRender.com. (B and C) Imino regions of 1D ^1^H-NMR spectra of s2m-Cov-2 (B) and s2m-CoV-1 (C) annealed in the presence (red) or absence of Mg (black) (25 mM Na Phos, 50 mM NaCl, pH 6, 298 K).

Here, we utilise a synergistic approach that combines NMR spectroscopy with CLIR-MS, a method that couples cross-linking and mass spectrometry (MS) ^17,18^ to characterise the interactions of the two structured RBDs of the N-protein with a folded RNA element, towards the final goal to identify interaction hotspots and inform a drug screening campaign (Figure 1A). As a proxy of structured viral RNA, we use s2m, a well folded and conserved RNA element that is located at the 3’UTR region and that is known to interact with the N-protein^19,20^. By combining the cross-linking sites with NMR chemical shift perturbations, we identified the interface of binding of both domains to s2m and used this information for guiding an NMR fragment screening by selecting molecules binding at these interfaces. The identified molecular fragments are promising modulators of N-protein RNA interactions that can be used as starting point for further drug development.

## MATERIALS AND METHODS

### Protein expression

Individual domains were expressed and purified as previously reported^21^. All nucleocapsid protein constructs were expressed in Escherichia coli BL21 (DE3) overnight at 18°C after induction at an optical density of 0.6 with 0.6 mL of 1 mM isopropyl-β-D-1-thiogalactopyranoside (IPTG). Cells were harvested by centrifuging at 3500 g and resuspended in 20 mM Tris (pH 8.0) and 1 M NaCl buffer; lysed by a cell cracker, and centrifuged again at 34960 g at 4°C. The supernatant was passed through Ni-NTA agarose resin (Quiagen) to immobilise the polyhistidine-tagged domains. Proteins were eluted with 20 mM Tris (pH 8), 500 mM NaCl, and 300 mM imidazole. Samples were then dialyzed against 20 mM Tris (pH 8), 300 mM NaCl, and 5 mM 2-mercaptoethanol at 4°C overnight. Following TEV cleavage (in-house purified) and removal of the excess N-terminal tag and TEV by Ni affinity, samples were additionally subjected to size exclusion chromatography (SEC; Superdex 75/200) into the NMR buffer (25 mM Na-phosphate pH 6.0, 50 mM NaCl).

### RNA in vitro transcription

RNAs were prepared from synthetic DNA templates containing the T7 promoter region. The last two nucleotides at the 5’ end of the template strand were modified to contain 2’OMe groups. The respective DNA templates were annealed with a shorter DNA strand containing the T7 promoter region plus one G nucleotide. These synthetic DNAs were purchased from Microsynth, Switzerland. The in vitro transcription reaction (using in-house purified T7 polymerase) was optimized by varying DMSO, MgCl_2_ and NTPs concentration. Optimized in vitro transcription reactions were incubated for at least 5 h at 37 °C and supplemented with EDTA before anion exchange HPLC purification using a DNAPac-PA100 22x 250 mm column heated to 85°C. After column equilibration and injection at 100% buffer A (6 M urea, 12.5 mM TRIS-HCl, pH 7.4), a 0.25 min gradient to 15% buffer B (6 M urea, 12.5 mM TRIS-HCl, pH 7.4, 0.5 M NaClO_4_) was used to elute excess of mononucleotides. The RNA was eluted using a slower gradient from 15% to 73% buffer B over 12.5 min, and the column was washed to 100 % buffer B and re-equilibrated to 100% buffer A before the following injection. The pure RNA fractions were identified by urea-PAGE, the RNAs isolated by butanol extraction and lyophilized before resuspension into the NMR buffer (25 mM Na phosphate pH 6, 50 mM NaCl). The RNA constructs were then annealed in the presence or absence of MgCl_2_ by heating to 98°C for 1 min and snap-cooled on ice for 10 minutes before use.

### Cross-linking of stable isotope labelled RNA coupled to mass spectrometry (CLIR-MS)

#### UV cross-linking

The CLIR-MS experiments were performed as previously described^22^. Purified nucleocapsid protein samples were first mixed with the viral s2m element RNA in a protein:RNA ratio ranging between 1:1 to 2.5:1 followed by UV cross-linking at 254 nm. In detail, all mixtures were split into aliquots of 20 µL each and spotted on a 60-well microtitre plate (Sarstedt). The plate was placed on ice at a distance of 2 cm to the UV lamps in a Spectrolinker XL-1500 UV Crosslinker (Spectronics Corporation) at a total irradiation energy of 4.8 J/cm^2^. The distance to UV lamps was maintained constant by using a 3D-printed ice bucket holding the 60-well plate. Then, the cross-linked samples were recovered and mixed with 1/10 volume of 3 M sodium acetate (pH 5.2) and 5 volumes of ice-cold ethanol. The precipitated samples were stored at -20 ^°^C for at least 12 h. Next, the cross-linked samples were centrifuged at 4 °C for 30 min at 16,000 × g followed by washing the pellet with 80% ethanol. A second centrifugation step was performed as above, the supernatants were removed and the pellets were dried at room temperature for 10 min. The dried samples were resuspended in 50 mM Tris-HCl (pH 7.9) and 4 M urea, followed by a dilution to 1 M urea with 50 mM Tris-HCl (pH 7.9) prior to RNA digestion. The RNA digestion was performed by addition of 5 U of RNase T1 (Thermo Fisher Scientific) and 5 μg of RNase A (Roche) per mg of cross-linked complex and incubated at 52 °C for 2 h. The protein was digested at 37 ^°^C overnight by adding sequencing-grade trypsin (Promega) at a 24:1 substrate-to-enzyme ratio. Trypsin was inactivated by increasing the temperature to 70 ^°^C for 10 min. After protein and RNA digestions, samples were cleaned-up by solid-phase extraction using Waters C18 SepPak columns (50 mg), followed by a drying step under vacuum. The dried samples were then isotope labelled as previously described^22^. In detail, samples were dissolved in 86 μL of water, 10 μL of 10x T4-PNK buffer (NEB), 1 μL of 100 mM ATP, 1 μL of 100 mM ^18^O_4_-γ-ATP followed by addition of 20 U of T4-PNK (NEB) and incubation for 1 h at 37 °C. Another solid-phase extraction step was performed as above. Peptide-RNA conjugates were enriched afterwards by using TiO_2_ beads. First, 5 mg of TiO_2_ beads (Titansphere PhosTiO 10 μm, GL Sciences) were equilibrated with 500 μL of loading buffer (50% acetonitrile, 10 mg/mL lactic acid, and 0.1% trifluoroacetic acid (TFA)) for 15 min. Then, the dried samples were dissolved in 100 μL of the loading buffer and incubated for 30 min with the already equilibrated beads. After a centrifugation step and removal of the supernatant, the beads were washed sequentially with 100 μL of the loading and washing buffer (50% acetonitrile and 0.1% TFA). The cross-linked peptides were eluted twice with 75 μL of the elution buffer (50 mM (NH_4_)_2_HPO_4_ (pH 10.5)), and the solution was immediately acidified by adding 10 μL of TFA. Cross-linked peptides were further purified by StageTip solid-phase extraction. Briefly, two layers of C_18_ membranes (3M Empore) were first washed with (1) 70 μL 100% acetonitrile (ACN), (2) 70 μL 80% ACN with 0.1% formic acid (FA), and (3) two times with 70 μL 5% ACN with 0.1% FA. Then, the samples were loaded and the tips were washed three times using 70 μL 5% ACN with 0.1% FA and finally eluted three times using 50 μL 50% ACN with 0.1% FA. The eluate was collected in LoBind tubes (Eppendorf), and samples were evaporated to dryness under vacuum.

#### Liquid Chromatography−Tandem Mass Spectrometry

The dried samples were resuspended in 20 μL of water/acetonitrile/formic acid (95:5:0.1, v/v/v) and 5 μL were analysed by LC-MS/MS on an Orbitrap Fusion Lumos mass spectrometer (ThermoFisher Scientific) equipped with a Nanoflex electrospray source coupled to an Easy nLC 1200 HPLC system (ThermoFisher Scientific). The cross-linked peptides were separated on a PepMap RSLC column (250 mm × 75 μm, 2 μm particle size, ThermoFisher Scientific). Chromatographic separation was performed with a 60 min gradient starting at 6% and increasing to 40% B (mobile phase A: water/acetonitrile/formic acid (98:2:0.15, v/v/v); mobile phase B: acetonitrile/water/formic acid (80:20:0.15, v/v/v)), with a flow rate at 300 nL/min. The Orbitrap Fusion Lumos was operated in positive ion, data-dependent acquisition mode. Acquisition was performed at a resolution of 120,000 in 3 s cycles. During each cycle, precursor ions were selected for fragmentation using stepped higher energy collision-induced dissociation (normalized collision energy, 23 ± 5%). Fragment ions were detected in the Orbitrap at a resolution of 30,000, with an isolation window of 1.2 *m/z*, a dynamic exclusion duration of 30 s and selected charge states = 2-7^+^.

#### Data Analysis

Mass spectrometry data generated in Thermo .raw format were converted into mzXML format using msconvert ^23^ and analysed using a modified version of xQuest^17,24–26^. The searches used a database containing the target protein sequence and its reversed sequence. Each amino acid was considered as a possible cross-linked site and nucleotide adducts of up to four residues were allowed. Combination of different neutral losses were specified based on the findings described elsewhere^24^. Main xQuest search parameters were as follows: isotope mass shift for post-digest labelling of 6.012735 Da, mass tolerance: 15 ppm, retention time tolerance for matching heavy and light spectra: 60 s, enzyme used: trypsin, maximum number of missed cleavages: 2, MS^1^ mass tolerance: 10 ppm, MS^2^ mass tolerance 20 ppm. A 1% false discovery rate cut-off at the unique peptide-RNA conjugate level was used to control the error rate for identifications.

### NMR spectroscopy and library screening

All NMR spectroscopy measurements were performed using Bruker AVNEO 500 MHz, AVIIIHD 600 MHz and AVNEO 600 MHz spectrometers equipped with cryoprobes. 1D ^1^H spectra were acquired in 10% D_2_O using WATERGATE for water suppression, with a typical spectral width of 16.6 ppm. 1D SOFAST experiments centered on the imino region (12.5 ppm) were acquired with 0.2 s interscan delay^27^. Typically, 2D [^15^N, ^1^H] HSQC spectra were acquired with spectral widths equivalent to 16 ppm for ^1^H and 36 ppm for ^15^N centered at 4.699 ppm and 117 ppm, respectively. 1D ^19^F NMR experiments were performed with a spectral width of 50 ppm centered at a frequency corresponding to -70 ppm for CF_3_ containing molecules, while a spectral width of 150 ppm centered at -120 ppm was used for the remaining fluorine containing fragments, which were also subjected to ^1^H-decoupling to prevent splitting and therefore loss of signal intensity. Ligand binding was monitored by measuring ^19^F T2 relaxation with two relaxation periods (typically 5–10 ms and 200–300 ms) in the presence or absence of the macromolecules. Mixtures of fragments were prepared to contain 24–30 molecules with not overlapping ^19^F chemical shifts. The final concentration of each fragment in the mixture was 40 µM and the final macromolecule concentration was varied between 5 and 10 µM. The data were processed using Topspin 4.1 (Bruker) and analyzed with NMR-FAM-SPARKY^28^. Protein and RNA assignments were transferred from BMRB deposition entries 34511 (NTD), 50518 (CTD), and 50341 (s2m). Combined ^1^H and ^15^N chemical shift perturbations were calculated using the following equation:

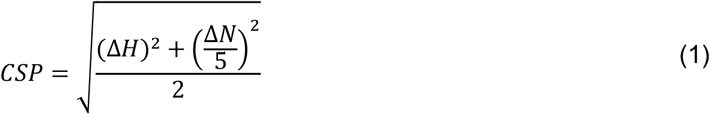

Estimated K_D_ values were calculated by fitting the chemical shifts of selected amino acid residues upon ligand titration according to the following equation in Origin:

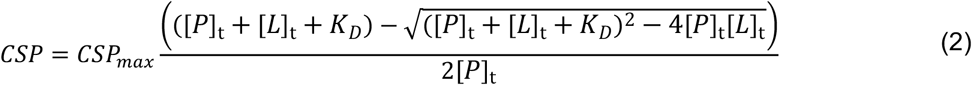

Where [P]_t_ is the total protein concentration, which was approximated to be constant; and [L]_t_ is the ligand concentration.

### Molecular modelling and small molecule docking

The RNA model of s2m-CoV-2 was generated from the RNAcomposer websever by restraining base pairing according to the experimental observation of the imino protons^29,30^. Structures of CTD and NTD were obtained from the PDB 6YUN and 6YI3. After pdb preparation, the macromolecules were docked according to a standard HADDOCK protocol adapted for nucleic acid docking in the HADDOCK 2.4 webserver using the guru mode^31,32^. NTD and CTD residues with perturbation larger than 2-fold standard deviation upon s2m-CoV-2 addition were used as ambiguous NMR restraints. CLIR-MS restraints were used in the following way: high-count cross-links with ambiguous nucleotides sequences were treated as ambiguous restraints; high-count cross-links coupled with unambiguous nucleotides were used as unambiguous 12 Å upper limit distance restraints between N1/N9 atoms of the nucleotides and Cα of the amino acid residue. Overall, better scoring models were also in agreement with experimental observation.

Small molecule docking was also performed in HADDOCK using the optimised protocol for ligand docking. Small molecule coordinates were generated starting from sdf files, which were converted to 3D structures using openbabel^33^. Protein residues with amide chemical shift perturbations larger than 2-fold the standard deviation were used as ambiguous restraints. The entire ligand was treated as active residue in the docking protocol. Also in this case, better scoring models were in general agreement with the experimental observation. Graphical representations of the models were generated using ChimeraX^34^.

## RESULTS

### The RNA s2m-CoV-2 is a stable folded element

Before utilising our hybrid structural approach to elucidate protein-RNA interactions, we relied on NMR spectroscopy to confirm the folding of the RNA elements to be used in our study. We choose the stem-loop 2 (s2m) located at the 3’UTR of the SARS-CoV-2 genome because the interaction of the N-protein with this part of the viral RNA was previously reported in the context of SARS-CoV-1. Furthermore, the structure of a single nucleotide variant of the current s2m-CoV-2 (G33U) has been determined by x-ray crystallography revealing a stable but unusual RNA folding mediated by two atoms of magnesium^35^. S2m is a 43 nucleotides (nt) RNA element that is conserved in a variety of virus species, including *Coronaviridae, Astroviridae, Caliciviridae* and *Picornaviridae*^36^, but its function is still unclear. However, in vitro and in vivo RNA structural analysis, in vivo protein cross-linking, and antisense oligonucleotide targeting have confirmed that s2m is folded in the context of the entire genome, and has a role in the regulation of the viral cycle^20,37,38^. Importantly, s2m of SARS-CoV-2 has a two nucleotides modification relative to SARS-CoV-1 - U7C and G33U (Figure 1A).

In our study, we systematically adapted the natural RNA constructs at the 3’ and 5’ termini by replacing the two terminal unpaired nucleotides with two G-C base pairs in order to facilitate in vitro transcription, as previously reported^20^. At first, we compared s2m-CoV-2 and s2m-CoV-1 by measuring NMR of the imino protons by ^1^H NMR spectroscopy. This provides rapid information on base pairing and therefore RNA folding in solution (Figure 1B and 1C). Consistent with previous studies, s2m-CoV-2 is well folded in the absence of magnesium. Furthermore, this adopts a secondary structure that deviates substantially from a previous x-ray crystal structure of the single-nucleotide mutant^20^. Under the same conditions, the imino signals of s2m-CoV-1 are substantially reduced in number and intensity, suggesting a more dynamic and/or unfolded structure. Refolding the RNA in the presence of 5 mM MgCl_2_ resulted in a stabilisation of the secondary structure as suggested by sharper and additional imino resonances. This effect was not observed with s2m-CoV-2, which upon refolding in the presence of MgCl_2_ retained the same overall conformation even though imino protons experience line broadening (due to conformational exchange or increased relaxation associated with the presence of traces of manganese). Collectively, these data suggest that the two-nucleotide substitution present in s2m-CoV-2 stabilises the RNA structure relative to s2m-CoV-1. Based on this information, a 3D model of s2m-CoV-2 was generated with the RNAcomposer websever^29,30^ to be used in our subsequent studies.

### s2m folding favours specific interactions with the CTD

As first step of our investigation, we studied the interaction between s2m-CoV-2 and the CTD. As previously reported, the CTD is a dimer in solution, of which each monomer is composed of three 3_10_ helices, five α-helices and two β-strands; these last two are intertwined in an antiparallel fashion to form the dimerization interface^39^. The s2m-CTD interaction was previously reported for SARS-CoV-1 and more recently examined in the context of SARS-CoV-2, however, by using a DNA analogue of s2m-CoV-2, which is unlikely to recapitulate the RNA secondary structure and 3D shape^40,41^. Initially, we performed NMR titrations to observe the RNA imino resonances upon addition of the CTD. Figure 2A shows small (1–6.5 *Hz*) chemical shift perturbations that were induced upon addition of increasing amounts of the CTD. Chemical shift perturbations were localised around the residues of the two helices surrounding the asymmetric internal loop of s2m-CoV-2 (G30–G34 and G41–U3). At the same time the RNA signals were broadened significantly and at two equivalents of CTD the signals were hardly visible. Figure 2B further illustrates chemical shift perturbations of the amide backbone of the CTD upon RNA binding that was monitored by ^1^H-^15^N HSQC upon titration of s2m-CoV-2 to a ^15^N-labelled CTD. The binding event is in the intermediate exchange regime on the NMR time scale, which resulted in almost complete signal loss already upon addition of 0.1 equivalent of s2m-CoV-2. This prevented an exact mapping of the residues involved in the interaction at a stoichiometric ratio of s2m-CoV-2 and CTD. However, we could gather some insights into the CTD residues involved in the interaction by adding small aliquots of s2m-CoV-2 (0.02-0.03 equivalents relative to the CTD). Under these conditions, it was possible to observe relatively large chemical shift perturbations considering the small amount of RNA added. The residues at the N-terminus of CTD were particularly easy to follow as their signal decays more slowly because of the intrinsically dynamic nature of this N-terminal region (Figure 2D, S1A). This flexible helical region is rich in basic amino acid residues and exhibited the largest chemical shift changes (assigned peaks in Figure 2B) in agreement with earlier findings that this region is essential for RNA binding in the context of SARS-CoV-1^41^. Other small chemical shift perturbations were observed for sparse residues of the CTD, but signal broadening at higher RNA ratio prevented the unambiguous mapping of any other sites of interaction (Figure 2D).

**Figure 2.**
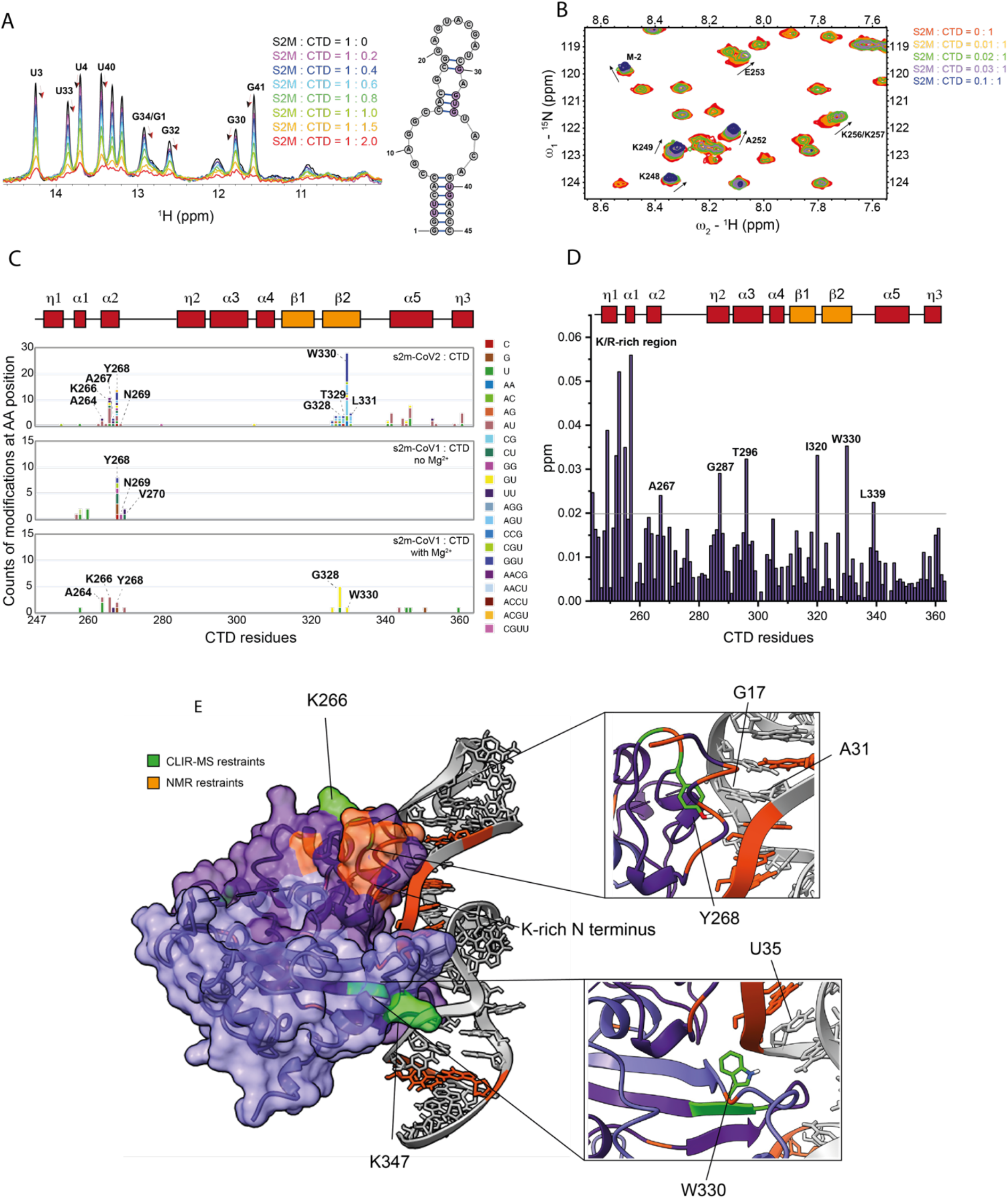
(A) Overlay of 1D-^1^H-SOFAST NMR spectra focused on the imino resonances of s2m-SARS-CoV-2 (50 *µ*M) at increasing ratios of CTD (25 mM Na Phos, 50 mM NaCl pH 6, 298K). The arrows highlight the chemical shift perturbation of selected residues. (B) Overlay of ^1^H-^15^N HSQC spectra highlighting the chemical shift perturbations of selected CTD (100*µ*M) residues upon addition of small aliquots for s2m-CoV-2 (25 mM Na Phos, 50 mM NaCl pH 6, 303 K). (C) Depiction of the cross-linking counts obtained by CLIR-MS for each amino acid residue of CTD upon addition of s2m-CoV-2, s2m-CoV-1 without Mg^2+^ and s2m-CoV-1 in the presence of Mg^2+^. (D) Plot of combined ^1^H and ^15^N chemical shift perturbation induced by s2m-CoV-2 to the CTD amino acid residues at 0.03:1 ratio before complete signal broadening. (E) HADDOCK model 1 of the interaction between CTD and s2m-CoV-2 with highlighted experimental restraints used and focus on key interaction sites.

For this reason, we complemented the NMR-derived information by performing CLIR-MS experiments. This method identifies UV protein-RNA cross-linked products by MS and defines the amino acid and nucleotide residues that are in close proximity upon interaction^17^. Using CLIR-MS, we identified two CTD regions that are contacting RNA (Figure 2C), which became apparent when CTD was in 2-fold excess relative to s2m-CoV-2 (1:1 ratio CTD dimer to RNA). The most abundant interactions were observed for amino acids T329, W330 and L331 located in β-strand 2 at the dimerization interface, and residues K266 and Y268, in close spatial proximity to the basic region identified by NMR. These sites are also consistent with the chemical shift perturbations for residues A267 and W330 that were observed at 0.03:1 s2m-CoV-2:CTD ratio in the NMR experiments. It is noteworthy that the basic helical region at the N terminus that was identified as CTD site of interaction by NMR was not detected by the CLIR-MS experiment. This was expected as the lysine- and arginine-rich CTD sequence is digested by trypsin into small di- and tri-peptides, and detection by mass spectrometry is therefore challenging. While these positively-charged residues were expected to interact with RNA, the identification of the site at the β-strand 2 was surprising as it is located opposite to the positively-charged groove formed by the CTD dimer, which was deemed the more likely RNA binding site due to its electrostatic properties and shallow topology^11^. A more detailed interpretation of the distribution of RNA adducts on the amino acid sequence suggested that residue W330 is unambiguously located in proximity of the central asymmetric loop of s2m-CoV-2 as indicated by cross-links with trinucleotides GGU, AGU and CCG and the dinucleotides AC and CG that are present on both sides of the unpaired region. Cross-linking counts of the neighbouring T329 and L331 were enriched for trinucleotides GGU and AGU and dinucleotides CG and AC, which suggested that these residues are in proximity of the GUAC tetranucleotide at the 3’ side of the asymmetric loop. As a result of the higher degree of cross-linking observed at G and U residues, we inferred that the peptide T329-W330-L331 should be spatially close to the G34-U35-A36 stretch of s2m-CoV-2. This result also corroborated with NMR analysis that indicated the region surrounding the internal loop as the binding interface.

Analysis of the cross-linking counts of residue K266 and Y268 prevented unambiguous mapping of interactions, as nucleotides stretches that are distant in the s2m-CoV-2 construct were equally present, which might indicate multiple possible conformations. Further investigation of the RNA binding mode of the CTD was performed by CLIR-MS using the two-nucleotide mutant s2m-CoV-1 under the same conditions (in which s2m-CoV-1 should be only partially folded) or in the presence of 5 mM MgCl_2_, with the aim of pinpointing any effect deriving from the RNA sequence and shape (Figure 2C). Y268 and neighbouring residues corresponding to the positively charged region of the CTD cross-linked with relatively high counts under both conditions, while W330 only cross-linked when the s2m-CoV-1 was properly folded in the presence of magnesium. In this case, W330 higher crosslinking at the GU dinucleotide instead of the trinucleotide GGU was observed for s2m-CoV-2. We surmise that this is due to reflect the nucleotide mutation U33-G33 of s2m-CoV1, corroborating again the indication that the dimerization interface of CTD interacts with the loop region of s2m. The cross-linking pattern of s2m-CoV-2 in the presence of Mg was also similar to that observed without Mg^2+^, confirming that the overall shape of s2m-CoV-2 is independent of [Mg^2+^] (Figure S1D). Overall, these data suggest that the site at the dimerization interface interacts preferentially with folded RNA motifs and might indicate some shape-specific recognition mode of the CTD towards certain RNA elements.

### The CTD dimer binds to s2m-CoV-2 via a cleft at the dimerization interface

Next, we used the NMR and CLIR-MS sparse restraints obtained in the absence of Mg to construct a model of the CTD-s2m-CoV-2 interaction by using HADDOCK^31,32^. The two conformations with the best docking score were consistent with our experimental data and suggest that the CTD binds to s2m-CoV-2 as a dimer (Figure 2E and Figure S1C). These two conformations are similar, however, with some relevant differences such as the orientation of the s2m-CoV-2 double-helix, which is rotated by 180 degree, potentially explaining the ambiguity of the interaction site of s2m-CoV-2 with residue Y268 (Figure S1C). In both models, W330 at the dimerization interface of the CTD is in contact with the major groove side adjacent to the asymmetric internal loop of s2m-CoV-2. However, model 1 seems somewhat more plausible as W330 is in proximity of residue U35 as expected (Figure 2E). In model 2, W330 is instead close to the opposite side of the asymmetric loop near residue C7 (Figure S1C). The positively-charged helical region at the N-terminus of the CTD, which includes the residues identified by NMR and Y268 is placed in contact with the minor groove above the internal loop in model 1. Y268 is in proximity of the unpaired residue A31, which is consistent with the cross-link with the tetranucleotide -ACGU-detected by CLIR-MS (Figure 2D and 2E). In contrast, in model 2, this region is instead placed below the asymmetric loop, with Y268 next to residue C44, this time accounting for the -AC-dinucleotide observed in CLIR-MS. Further validation of the models could also be gathered from residue K347 in α-helix 5, despite the lower cross-linking counts in the CLIR-MS experiments. Indeed, the contact of this residue with RNA strongly suggests that the CTD is binding in a dimeric form since the identified main binding interface (i.e., W330) and K347 within same monomer are located at opposite sides of the domain. Cross-linking of residues K347 are enriched in GU and AU nucleotides in agreement with model 2 that places K347 close to G34-U35-A36, but also to model 1 if one considers that a cross-link to AU might also indicate a cross-link to AC due to photo-mediated deamination of C to U (in model 1, K347 is in close proximity to A6-C7)^42^. A recent study revealed that π-π stacking interactions between aromatic amino acid residues and nucleobases are the most dominant type of interactions for cross-linking to take place^42^. Our models align well with this observation with the majority of cross-linking events associated with the proximity between aromatic amino acid residues (i.e., Y268, W330 and F346) with flexible unpaired regions of the RNA. This biased behaviour can also explain why the counts of cross-links related to the different conformations are not proportionally distributed and cannot be used to estimate the relative ratio of the species. To further validate our model, we also docked the CTD to the x-ray determined construct (PDB 1XJR), the structure of which predominately differs from our model in the central loop region. The best scoring conformations recapitulate the same binding mode with the β-sheet region of the CTD inserting into the major groove and the helical basic region in close proximity to the minor groove of s2m (Figure S1E).

By investigating the overall structure of the complexes, we conclude that only a small portion of the large positively charged groove of CTD is occupied by s2m-CoV-2 (Figure S1B). This is unexpected as this positive region was anticipated to be the main RNA binding site^41^, but confirms a recent study that identified the cleft around residue W330 as a guanine binding pocket^43^. For this reason, we hypothesize that a second binding event on the opposite side of the CTD as well as other alternative RNA binding modes are required for genome packing. Taken collectively, we have identified the dimerization interface of CTD and the central region of s2m-CoV-2 as potentially specific druggable hotspots to prevent RNA binding.

### NTD interacts with s2m with high affinity and a distinct binding signature

As part of our workflow, we then applied the same approach to investigate the binding of s2m-CoV-2 to the NTD domain of the N-protein. The structure of NTD was previously solved both by NMR spectroscopy and x-ray crystallography and revealed a right hand-like fold comprising a β-sheet core formed by five antiparallel strands intertwined with two α-helices and a long basic β-hairpin, known to interact with nucleic acids^7,44^. In our analysis shown in Figure 3A, NTD-s2m-CoV-2 binding is in the fast-intermediate exchange regime on the NMR time scale, which resulted in larger and more easily observable chemical shift perturbations compared to the CTD, which allowed for mapping of both RNA bases and amino acid residues involved in the complex formation. Upon addition of up to three equivalents of the NTD to s2m-CoV-2, the imino protons G30-G32-U33 and G39-U40-G41 showed the largest chemical shift perturbations indicating that the central region of s2m-CoV-2 is the preferred binding site (Figure 3A). Titration of one equivalent of s2m-CoV-2 to the ^15^N-labelled NTD shown in Figure 3B and 3C identified the amino acid residues that are involved in the binding. These comprise regions that were previously identified as nucleic acid-binding such as the N terminus (i.e., G44, L45, and N47), the β-hairpin loop (K100, S105), and the β1-β2 strands of the β-sheet region. The overall chemical shift perturbations are also very similar to the one previously observed for dsRNA binding^7^.

**Figure 3.**
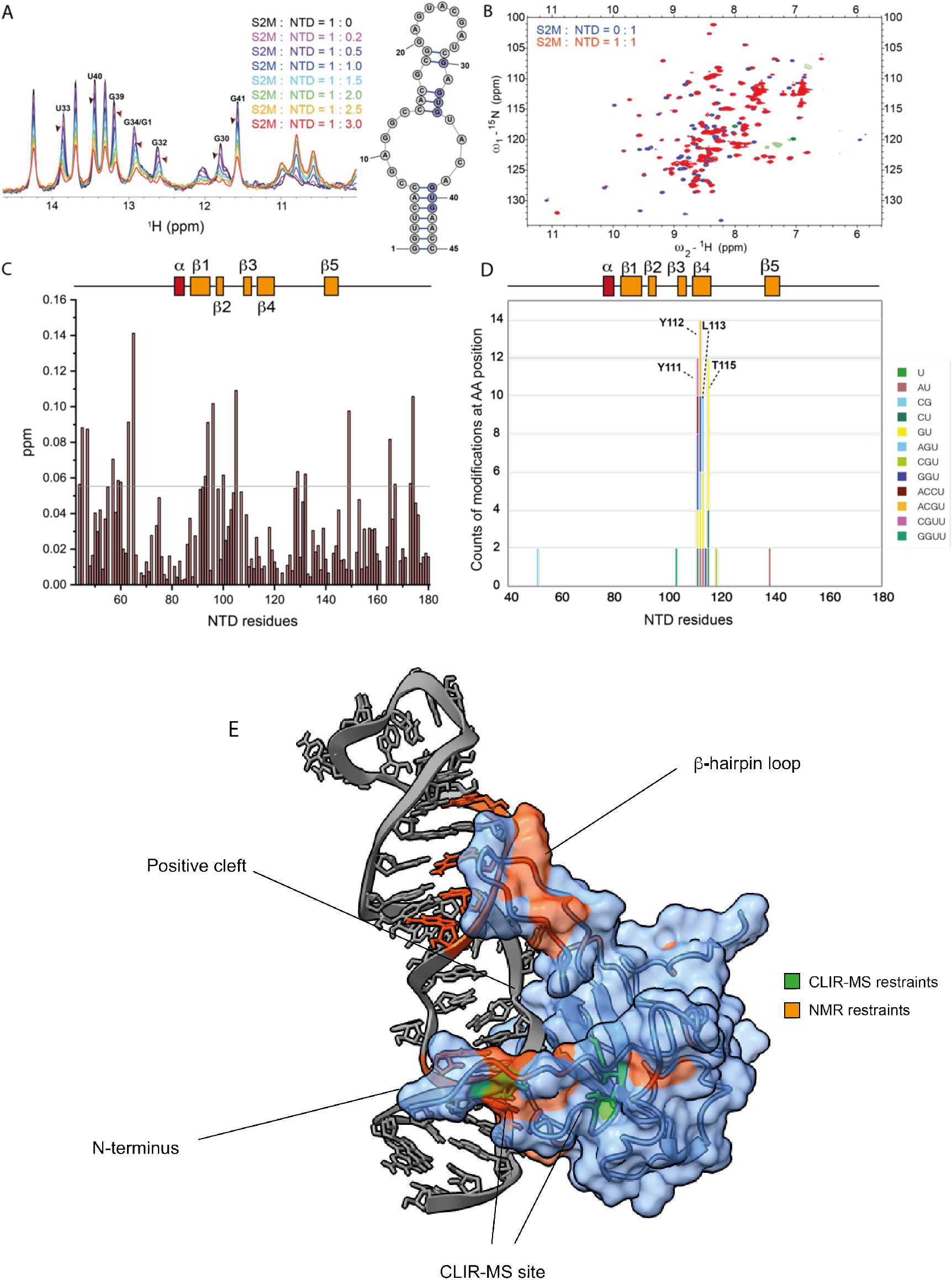
(A) Overlay of 1D-^1^H SOFAST NMR spectra focused on the imino resonances of s2m-SARS-CoV-2 (50 *µ*M) at increasing ratios of NTD (25 mM Na Phos, 50 mM NaCl pH 6, 298 K). The arrows highlight the chemical shift perturbation of selected residues. (B) Overlay of ^1^H-^15^N HSQC spectra of free NTD (100 *µ*M, purple) and NTD-s2m-CoV-2 at 1:1 ratio (100 *µ*M, red) (25mM Na Phos, 50 mM NaCl pH 6, 310 K). (C) Plot of combined ^1^H and ^15^N chemical shift perturbation induced by s2m-CoV-2 to NTD amino acid residues (the reference line refers to two standard deviations of the overall perturbations). (D) Depiction of the cross-linking counts obtained by CLIR-MS in relation to the amino acid residues of NTD when mixed with s2m-CoV-2 at ratio 2.5:1. (E) HADDOCK model of the interaction between NTD and s2m-CoV-2 with highlighted experimental restraints used.

Next, CLIR-MS analysis indicated the three amino acid stretch Y111-Y112-L113 and T115 were preferentially cross-linked. Residues Y112-L113 are located deep in the interior of the protein between β-strand 3 and 4 (Figure 3D) and we cannot exclude that the high cross-linking count is biased by a local redox potential favouring a radical transfer mechanism towards this peptide stretch^42^. This is possible also considering that Y112 is the only residue of the highly aromatic β-strand 3, which could be in close contact with the RNA. However, some degree of sequence-specificity is suggested by the identity of the cross-linked nucleotides, which include the trinucleotide -AGU-, -CUU- and the tetranucleotide -CGUU-, unambiguously placing this aromatic sequence to interact with the lower stem of s2m-CoV-2.

Analogously to CTD, we modelled the interaction between NTD and s2m-CoV-2 by combining CLIR-MS and NMR restraints with HADDOCK (Figure 3E). In this instance, the unambiguous modelling of the orientation of s2m-CoV-2 relative to the NTD was more challenging due to the lower crosslinking sites combined with the higher flexibility of the NTD relative to the CTD. However, our predicted mode of interaction recapitulated the binding mode previously reported for NTD towards dsRNA with some differences arising from the larger size and the central unpaired region of s2m-CoV-2 relative to the RNA used in other works^7^. Namely, the main contacts of the β-hairpin loop are with the backbone of the s2m-CoV-2 upper stem rather than with the major groove. The N terminus and the cross-linked peptide stretch are instead interacting with the minor groove side of the bottom stem of s2m-CoV-2. Analogously to the previous model of NTD binding to ssRNA^7^, the phosphate backbone of the central unpaired nucleotides of s2m-CoV-2 are interacting with the positively charged cleft of NTD.

Overall, our data confirm that the central cleft of NTD is mainly involved in RNA recognition and again the central region of s2m-CoV-2 is a potentially druggable hotspot to prevent this interaction^11^.

### A fragment screening identifies binders of s2m-CoV-2, NTD and CTD

As a further step of our approach to drug the interaction between the N-protein and s2m-CoV-2, we performed a fragment screening aimed at identifying binders of the three macromolecules (i.e., NTD, CTD and s2m-CoV-2). For each target, we used a fragment library composed of a total of 620 molecules containing at least one fluorine atom, namely 269 with a CF_3_ moiety and 351 containing a fluorine connected to an aromatic carbon. This library included fragments covering a diverse chemical space, with high solubility (> 300 µM in aqueous solution) and with suitable ^19^F NMR relaxation properties that allow for straightforward and fast identification of interactions^45,46^. In each NMR tube, each compound of a mixture of 24–30 molecules was screened at a 4–8 fold excess relative to the target macromolecule. The ^19^F T_2_ relaxation of the fragments in the presence of the target macromolecules was monitored and binders were identified when signal reduction was observed relative to the free molecules. Through this approach, we identified a total of 39 putative binders (16 s2m-CoV-2, 9 CTD and 14 NTD) that were confirmed by further NMR experiments with individual molecules. Fragments that showed binding at the interface of the RNA-N protein interaction were selected for further investigating their potential in disrupting the complex.

### CTD binders interact at the dimerization interface

Nine fragments showed increased relaxation in the presence of the dimerization domain CTD. ^1^H-^15^N HSQC protein observation mapping of the binding site of each compound in 14-fold excess relative to CTD (^15^N labelled) underscored two fragments (i.e., FL496 and FL232, Figure 4A, Figure S2) with clear chemical shift perturbations of residues that are in proximity to the dimerization interface. In particular, the peptide stretch of the β-2 strand comprising residues I333 and D337 was distinctively perturbed by FL496 as shown in Figure 4B. The similarity of FL496 with the purine nucleobases provides a further indirect confirmation that this CTD site has preference for nucleic acids^43^. These chemical shift perturbations were used as ambiguous restraints to dock the fragment to CTD that, as expected, placed FL496 at the RNA binding interface that was identified by CLIR-MS, offering potential for disrupting the interaction between CTD and s2m-CoV-2 (Figure 4D). Perturbation induced by the other compound FL232 were smaller but centred around the same putative binding site, comprising residues T329-W330-L331 and Q281-N285 (Figure S2B). However, ligand-observed experiments highlighted the promiscuity of both ligands, which also bind to s2m-CoV-2 as suggested by the increased ^19^F signal relaxation in the presence of RNA alone (Figure 4C, Figure S2C). The further signal broadening of the ligands in the presence of both s2m-CoV-2 and CTD suggested that both fragments are forming a ternary complex with CTD and s2m-CoV-2 (Figure 4C, Figure S2C). It is, however, impossible to conclude if in this complex the fragments are still bound to s2m-CoV-2, CTD or both due to the exchange regime of the CTD-RNA interaction that prevents epitope mapping of the complex. Overall, these data demonstrate that the site at the dimerization interface involved in RNA binding is indeed druggable and the identified bicyclic fragments, despite promiscuous, provide a starting scaffold for further chemical expansion.

**Figure 4.**
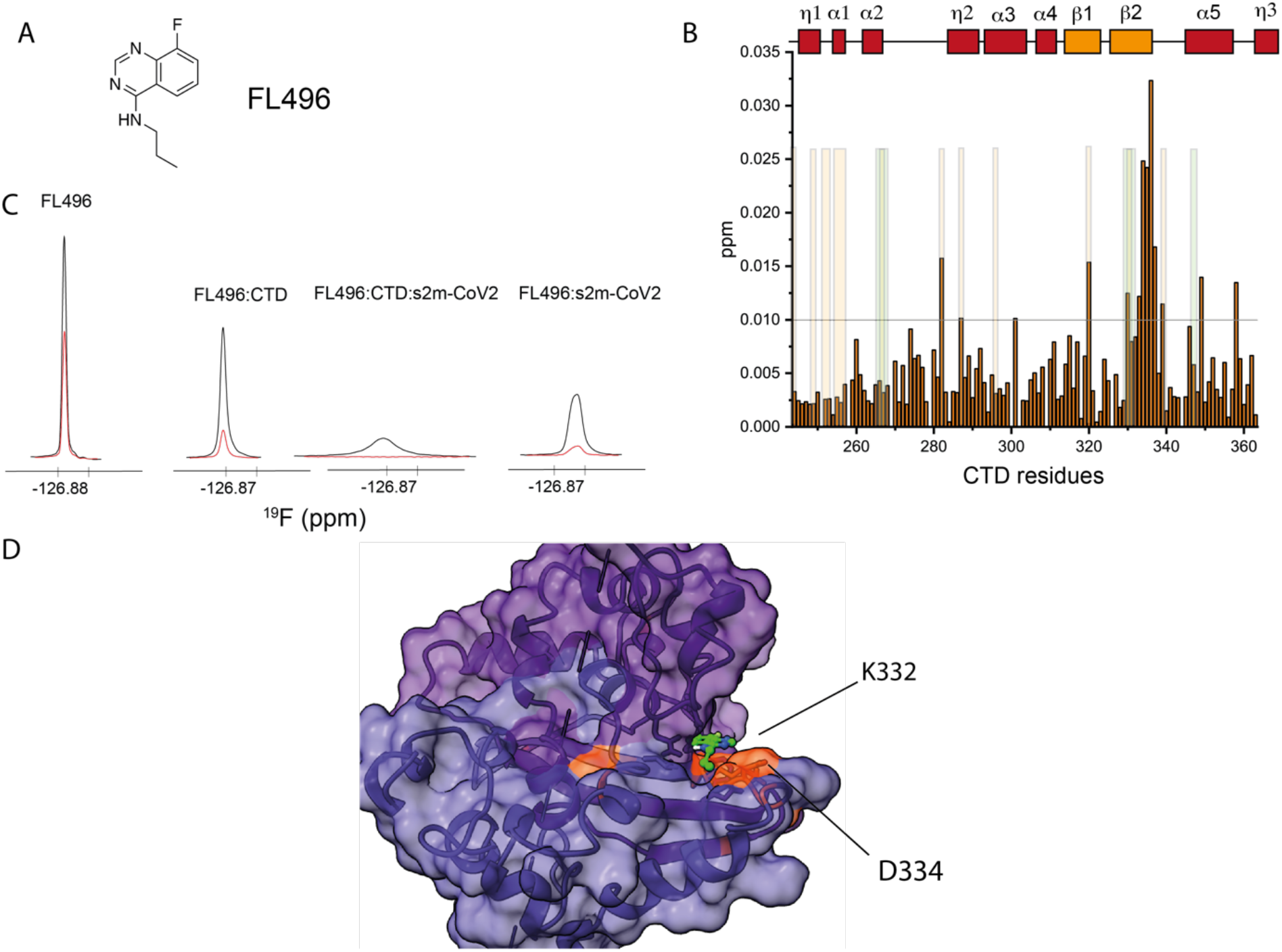
(A) Structure of the fragment FL496 identified as hit for CTD. (B) Plot of combined ^1^H and ^15^N chemical shift perturbation induced by FL496 (700 *µM*) to CTD (50 *µM*) amino acid residues (the reference line refers to two standard deviations of the overall perturbations; orange/green shaded areas serve as a reference to residues perturbed (NMR and CLIR-MS, respectively) by s2m-CoV-2 to facilitate the comparison between the effect of RNA vs FL496). (C) ^19^F resonances of FL496 in the presence or absence of CTD and s2m-CoV-2. Black: 1D ^19^F recorded using 10 ms relaxation delay; Red: 1D ^19^F recorded using 300 ms relaxation delay (CTD 50 *µ*M, fragment 750 *µ*M, RNA, 20 *µ*M, 25 mM Na Phos, 50 mM NaCl pH 6, 298 K) (D) HADDOCK model of the interaction between CTD and FL496 with highlighted key experimental restraints (orange).

### NTD binders act as potential disruptor of s2m interaction

The screening of NTD identified fourteen fragments with increased ^19^F relaxation, potentially indicative of binding (Table S4). This pool of fragments was enriched with a benzenesulfonamide scaffold (6 out of 14), suggesting a preferred chemical motif for NTD binding. ^1^H-^15^N HSQC spectra of each benzenesulfonamide compound in 15-fold excess relative to NTD (^15^N labelled) resulted in weak chemical shift perturbations mostly localised at β-strands 4 and 5 and the C-terminus (Figure S3A and S3B). While this weak effect prevented the unambiguous identification of the precise binding location, computational prediction coupled to the experimental observations never placed a representative molecule of the class (i.e., FL631) at the centre of the RNA binding interface (Figure S3D). The trifluoromethyladenine fragment ABL001 (Figure 5A) resulted in chemical shift changes for residues S51, R88, R89 and W108, which are located at the N-terminus and in β1 and β2 and form the RNA binding interface of the NTD (Figure 5B). Experimentally driven docking placed ABL001 in the pocket formed by these residues as highlighted in Figure 5D. This binding location is consistent with the AMP binding to the NTD of MERS-CoV that was previously reported^44^. The clear interaction between ABL001 and NTD is also corroborating our NTD-s2m-CoV-2 model, which shows direct contact between NTD and the unpaired bases of s2m-CoV-2. The dissociation constant (K_D_) was calculated in the low millimolar range (3–4 mM) by titrating ABL001 to the NTD in ^1^H-^15^N HSQC experiments (Figure S4). Despite the weak binding, we further investigated if this fragment had a potential disrupting effect on binding of the NTD to s2m RNA. Differently from the CTD binding fragments, the ^19^F relaxation of ABL001 in complex with NTD is only minimally increased in the presence of RNA. This suggests a competition between s2m-CoV-2 and ABL001, even though we cannot exclude that this small effect is due to the intrinsic properties of the fragment (Figure 5C). Furthermore, we found evidence that ABL001 affects the RNA binding by comparing ^1^H-^15^N HSQC spectra of the NTD-s2m-CoV-2 complex in the presence or absence of the ligand. In particular, the amide resonance relative to residue Y87 at the RNA binding interface is broadened by s2m-CoV-2, and appears as a sharp peak in the presence of ABL001 (Figure 5E). On the other hand, residues T49, T54, R95 and G96 are broadened in the presence of the ligand. Overall, this screening demonstrated that drugging the RNA binding interface of NTD is feasible by starting from analogues of the nucleic acid bases. Furthermore, we identified benzenesulfonamides as further interactors of NTD. These weak ligands are likely bound to a secondary pocket of NTD and chemical linkage of these two types of fragments might represent a potential avenue for increasing binding affinity and potency.

**Figure 5.**
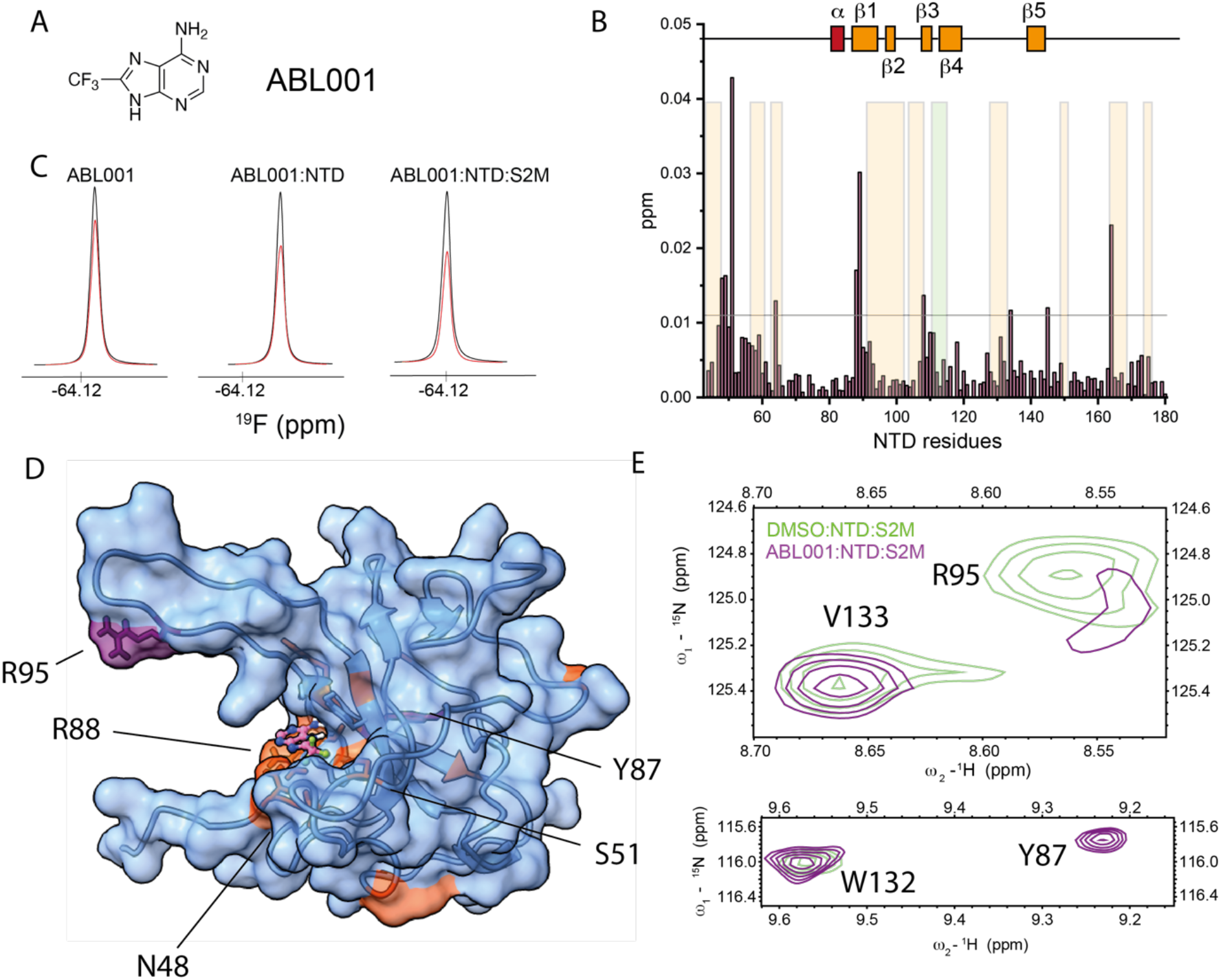
(A) Structure of ABL001 fragment identified as hit for NTD. (B) Plot of combined ^1^H and ^15^N chemical shift perturbation induced by ABL001 (1.5 mM) on NTD (50 *µM*) amino acid residues (the reference line refers to two standard deviations of the overall perturbations, orange/green shaded area serve as a reference to residues perturbed (NMR and CLIR-MS, respectively) by s2m-CoV-2 to facilitate the comparison between the effect of RNA vs ABL001). (C) ^19^F resonance of ABL001 in the presence or absence of NTD and s2m-CoV-2 indicating that ABL001 might compete with s2m-CoV-2 binding. Black: 1D ^19^F recorded using 5 ms relaxation delay; Red: 1D ^19^F recorded using 200 ms relaxation delay. (D) HADDOCK model of the interaction between NTD and ABL001 with highlighted experimental restraints used (orange). (E) Overlaid regions of ^1^H-^15^N HSQC of NTD-s2m-CoV-2 at 1:1 ratio (light green) and NTD-s2m-CoV-2-ABL001 (1.5 mM) (purple, 50 mM Na Phos, 100 mM NaCl pH 6.8, 310 K) highlighting the effect of ABL001 on selected residues at the RNA binding interface. In detail, R95 is broadened by the ABL001, while Y87 is sharpened. These residues are also highlighted in purple in (D).

### S2m-CoV-2 ligands compete for NTD binding

Finally, we also explored fragments binding to s2m-CoV-2 as means to disrupt the interaction with the N-protein binding domains. In total, sixteen fragments showed both increased ^19^F relaxation in the presence of s2m-CoV-2 and induced perturbation on the observable imino protons of the RNA (Figure 6A, Table S5). This screen identified common substructures suitable for further chemical refinement. For example, three molecules shared the [1,2,4]-triazolo-[4,3,b]-pyridazine core substituted at position 6 with an alkyl substituent connected to a positively charged moiety in the form of a cyclic tertiary/secondary amine, which is indicative of this chemotype binding to s2m-CoV-2 RNA (Table S5). Another subclass of fragment includes oxazole/oxadiazole rings connected through aromatic C-C bonds to fluorobenzene rings and further functionalised with positive charges in the form of primary, secondary and tertiary amines (Table S5). A similar scaffold was also identified in another screening campaign^47^.

**Figure 6.**
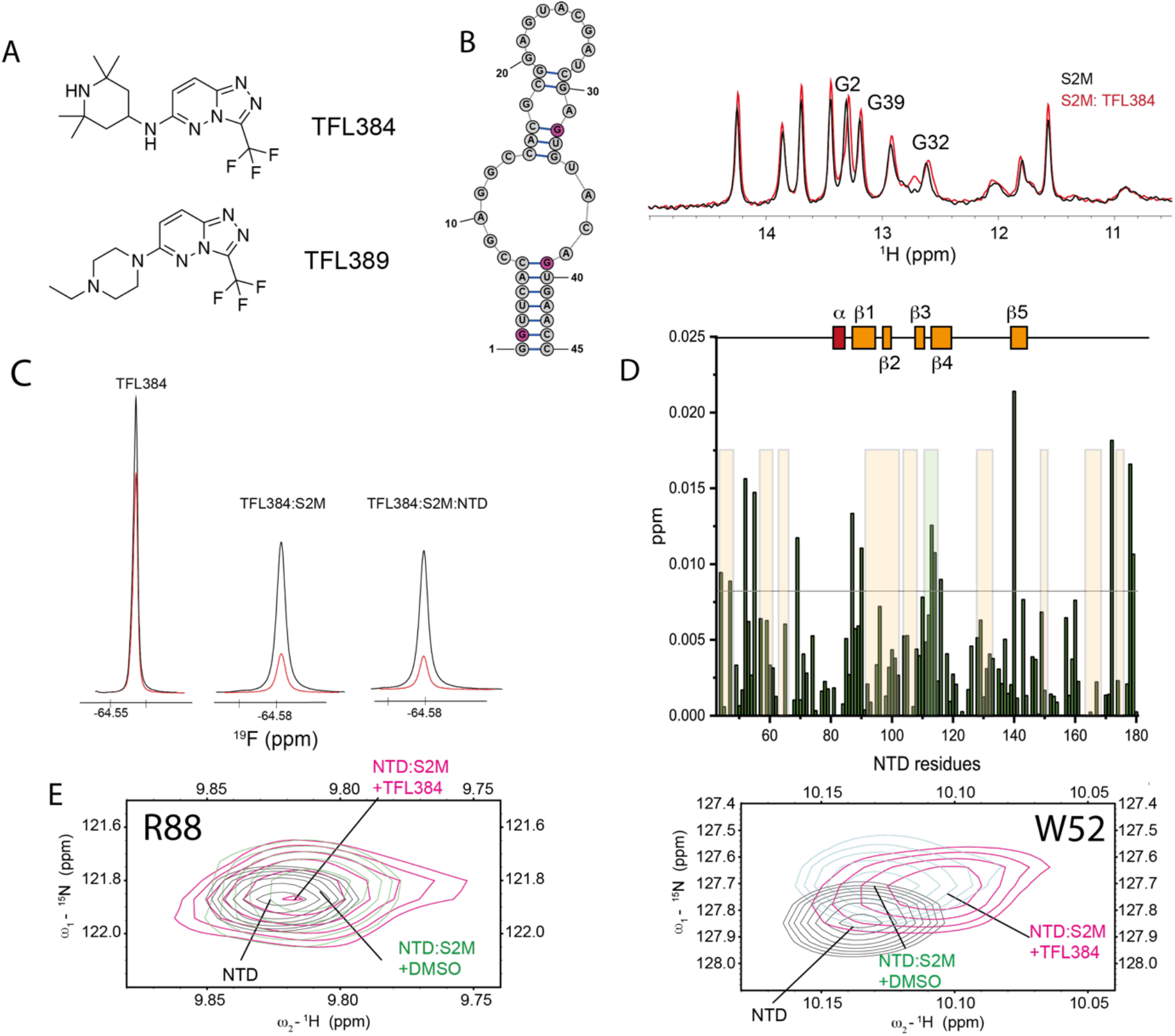
(A) Structure of fragments TFL384 and TFL389. (B) Overlay of 1D ^1^H-SOFAST NMR focusing on the imino region highlighting the chemical shift perturbation induced by TFL384 (1.5 mM) on s2m-CoV-2 (50 µM, 25 mM Na Phos, 50 mM NaCl pH 6, 298 K). (C) ^19^F relaxation of TFL384 in the absence and presence of s2m-CoV-2 and NTD. Black: 1D ^19^F recorded using 5 ms relaxation delay; Red: 1D ^19^F recorded using 200 ms relaxation delay (50 mM Na Phos, 100 mM NaCl pH 6.8, 310 K). (D) Plot of combined ^1^H and ^15^N chemical shift perturbations induced by TFL384 (1.5 mM) to NTD amino acid residues of the NTD-s2m-CoV-2 complex (100 µM; the reference line refers to two standard deviations of the overall perturbations, orange shaded areas serve as a reference to residues perturbed s2m-CoV-2 to facilitate the comparison between the effect of RNA in presence or absence of TFL384). (E) ^1^H-^15^N HSQC overlay of selected regions underscoring the effect that TFL384 has at the RNA binding interface of NTD (50 mM Na Phos, 100 mM NaCl pH 6.8, 310 K).

In this analysis, the RNA binding was monitored under two buffer conditions: buffer A (25 mM Na Phos, 50 mM NaCl, pH 6), which was optimal for imino NMR experiments, but more susceptible to artifacts at high fragment concentration; and buffer B (50 mM Na Phos, 100 mM NaCl, pH 6.8) which resulted in broadened imino signals with ambiguous assignment, but more robust towards high fragment concentration and still suitable for ligand and protein-observed experiments. The most frequent perturbations were observed for residues G2, G32, and G39 suggesting that the binders are sharing a similar binding site. The perturbed resonances belong to residues that are located at two distinct regions of s2m-CoV-2, both involved in binding with the N-protein. It is not clear if these shifts originate from multiple binding events of the fragments or from an allosteric effect upon binding. The binding monitored both by imino shifts and ^19^F relaxation was particularly clear for fragments containing the [1,2,4]-triazolo-[4,3,b]-pyridazine core, such as TFL384 and TFL389 (Figures 6B, 6C and Figure S5). We then investigated if selected fragments had any effect on CTD/NTD binding to s2m-CoV-2, by monitoring the signal of ligands (^19^F) and the change of the imino resonances upon addition of the protein domains (Figures 6C and D, Figures S5 and S6). The limited shift changes induced by the CTD on the imino resonances due to the intermediate exchange regime on the NMR time scale made the assessment challenging for this domain. On the other hand, we could observe some effects induced by the addition of the fragments when the NTD was added to s2m-CoV-2. This was more prominent for TFL389 that induced chemical shift changes of the imino resonances of U33, G2 and G39 relative to the DMSO control in buffer A (Figure S5). This effect was also recapitulated for both TFL384 and TFL389 in buffer B (Figure S6). Ligand observed experiments also indicated that the ^19^F relaxation of the ligand was only slightly increased upon addition of NTD to pre-formed s2m-CoV-2-TFL389 and s2m-CoV-2-TFL384 complexes suggesting that the fragments bind to s2m-CoV-2 in the presence of NTD and potentially affect NTD binding (Figure 6C and Figure S5). To confirm this, we monitored the amide backbone resonances of the s2m-CoV-2-NTD complex in the presence of the small molecules. Chemical shift perturbations of regions at the interface of s2m-CoV-2-NTD binding such as the N-terminus, β-1 and β-4 were observed for TFL384 and TFL389 (Figure 6D, Figure S5C). Evidence that s2m-CoV-2-NTD binding is affected was found in resonances corresponding to residues G69, G116, I74, R88, R89 and K143 that shifted towards the free form of NTD in the presence of TFL384. Other resonances (i.e., W52, A90, L113, F110, G129, Figure 6E), instead shifted in opposite direction. Overall, these data demonstrated that these weak RNA binders can represent the starting point for modulating the interaction between the NTD and s2m-CoV-2.

In order to inform further chemical derivatization, we also studied available analogues of TFL389. For example, the acetylated version of TFL389 (TFL383) resulted in reduced imino chemical shift perturbation and reduced effect on ^19^F relaxation suggesting that the positive charge of the piperidine is important for retaining RNA binding. Instead, removal of the trifluoromethyl group (TFL-2-194) did not preclude binding to RNA, suggesting that position 3 of the [1,2,4]-triazolo-[4,3,b]-pyridazine scaffold is a potential point for further chemical development (Figure S7).

## DISCUSSION

Here, we sought to find new starting points for drugs acting against SARS-CoV-2. In order to select the most promising candidates from fragment-based screening, we evaluated initial hits against structural information of the targeted protein-RNA complexes.

First, we characterized the interactions between the two RNA binding domains of the N-protein (i.e., NTD and CTD) and a structured RNA element involved in the regulation of the viral replication cycle (s2m-CoV-2). Our approach combines NMR and MS to determine the interface of interaction between RNAs and proteins, which in this case are involved in SARS-CoV-2 infection. One of the main advantages of combining NMR and CLIR-MS is the possibility to quickly obtain structural models from sparse restraints that reflect the solution behaviour of the macromolecules. While the level of structural details cannot be compared to high-resolution structural determination techniques, our approach is fast and overcomes limitations that arise from solution phenomena, such as liquid-liquid phase separation that results from protein-RNA interaction and might interfere with classical structural approaches. Importantly, our approach provides for the first time a model of interaction between the CTD and RNA. Our data suggest that the CTD binds to RNA in a dimeric form and in particular, CLIR-MS highlighted an unexpected interaction site at the dimerization interface that appears to be specific for structured RNA. This let us conclude that the CTD has multiple RNA binding modes as it recognizes RNA non-specifically through its large positively charged patch but might achieve some specific contacts through other interaction sites. Furthermore, our method expanded previous knowledge on how the NTD recognizes structured RNA elements of the viral genome.

In this paper, we demonstrated that this hybrid approach can rapidly provide candidates for drug development when combined with a fragment screening campaign. Targeting of protein-RNA interactions is indeed an emerging topic in drug discovery as the attention has shifted towards targeting RNA molecules and RNA binding motifs^48–51^. The N-protein of SARS-CoV-2 is a particularly attractive target as it is highly abundant during viral infections and has a fundamental role in regulating RNA packing. Here, we took advantage of the enhanced chemical diversity of a fluorinated fragment library to identify binders of the three macromolecules used in this work. The structural information gathered by the complementary NMR and CLIR-MS methods enabled us to filter fragments of interest and select molecules that have potential in disrupting protein-RNA interactions. We determined that the key complex formation hotspots of the NTD, CTD and s2m-CoV-2 are indeed targetable and that fragments can affect the protein-RNA binding, despite their expected weak binding affinity. Maybe unsurprisingly, the more promising fragments targeting NTD and CTD are predominantly bicyclic aromatic rings resembling the hydrogen bonding and shape of the natural nucleic acid bases. On the other hand, s2m-CoV-2 binders align well with known RNA-biased chemotypes^52^. These chemical scaffolds are suitable to further chemical functionalization to increase potency and provide well-characterized starting points for drug development for the treatment of COVID-19 or newly emerging SARS-CoV viruses.

## DATA AVAILABILITY

The mass spectrometry proteomics data have been deposited to the ProteomeXchange Consortium via the PRIDE^53^ partner repository with the dataset identifier PXD036671. NMR and modelling data have been deposited in Dryad (https://doi.org/10.5061/dryad.g4f4qrftb).

## SUPPLEMENTARY DATA

Supplementary Data are available at NAR online.

## ACKNOWLEDGEMENTS

We are grateful to the COVID-19 NMR consortium for organizing a global initiative to fight COVID-19 using NMR methodology and especially to Dr. Andreas Schlundt and Prof Martin Blackledge for providing plasmids for protein expression. GP thanks Prof Glenn A. Burley for his critical insight into a preliminary manuscript.

