## Supplementary Material for "A hybrid structure determination approach to investigate the druggability of the nucleocapsid protein of SARS-CoV-2"

### Supporting Information

#### 1. Supplementary Tables

**Table S1** Oligonucleotide sequences used for in vitro transcriptions and relative RNA products.

| | |
|---|---|
| **T7 promoter** | 5'-TAATACGACTCACTATAG |
| **Sm2-CoV-1 DNA template strand** | 5'O2′MeGO2′MeGTTCACTGTACCCTCGATCGTACTCCGCGTGGCCTCGATGAACCTATAG TGAGTCGTATTA |
| **Sm2-CoV-2 DNA template strand** | 5'O2′MeGO2′MeGTTCACTGTACACTCGATCGTACTCCGCGTGGCCTCGGTGAACCTATAG TGAGTCGTATTA |
| **S2m-CoV-1** | 5'GGUUCAUCGAGGCCACGCGGAGUACGAUCGAGGGUACAGUGAACC |
| **S2m-CoV-2** | 5'GGUUCACCGAGGCCACGCGGAGUACGAUCGAGUGUACAGUGAACC |

**Table S2** NTD and CTD amino acid primary sequence used in this work (numbering referenced to the UniProt entry P0DTC9 · NCAP_SARS2, in red amino acid residues that are not part of the natural sequence resulting from cleavage of the affinity tag).

| | |
|---|---|
| **NTD** | 44 51 61 71 81 91<br>GRLGLPNNTA SWFTALTQHG KEDLKFPRGQ GVPINTNSSP DDQIGYYRRA TRRIRGGDGK<br><br>100 110 120 130 140 150<br>MKDLSPRWYF YYLGTGPEAG LPYGANKDGI IWVATEGALN TPKDHIGTRN PANNAAIVLQ<br><br>160 170<br>LPQGTTLPKG FYAEGSRGGS |
| **CTD** | 247 253 263 273 283 293<br>GAMGTKKSAA EASKKPRQKR TATKAYNVTQ AFGRRGPEQT QGNFGDQELI RQGTDYKHWP<br><br>303 313 323 333 343 353<br>QIAQFAPSAS AFFGMSRIGM EVTPSGTWLT YTGAIKLDDK DPNFKDQVIL LNKHIDAYKT<br><br>FP |

**Table S3** CTD hits. Fragments that had enhanced $^{19}F$ relaxation in the initial screening.

| | | | |
|---|---|---|---|
| 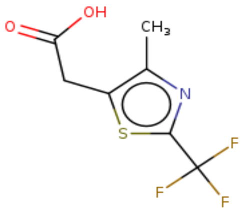 | BNC-TFL098 | 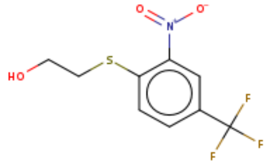 | BNC-TFL295 |
| 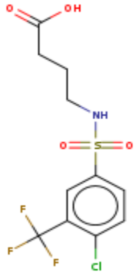 | BNC-TFL182 | 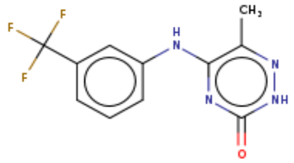 | BNC-TFL168 |
| 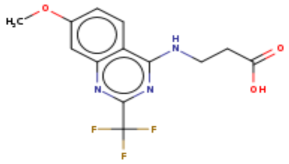 | BNC-TFL260 | 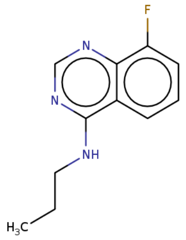 | BNC-FL0496 |
| 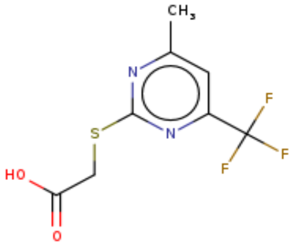 | BNC-TFL261 | 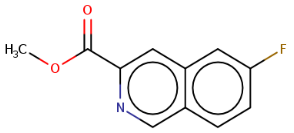 | BNC-FL0232 |
| 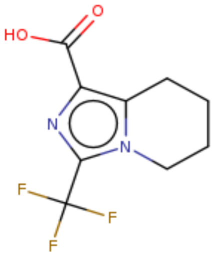 | BNC-TFL111 | | |

**Table S4** NTD hits. Fragments that had enhanced $^{19}$F relaxation in the initial screening.

| | | | |
|---|---|---|---|
| 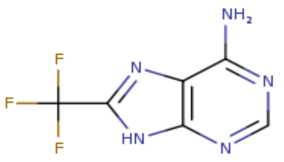 | BNC-ABL001 | 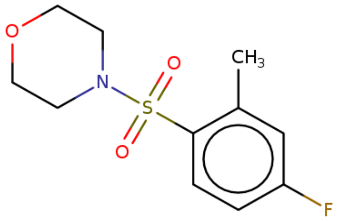 | BNC-FL0631 |
| 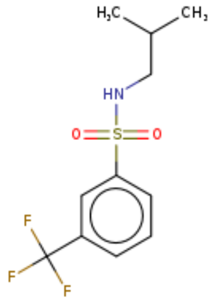 | BNC-TFL193 | 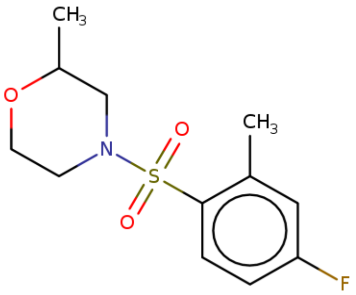 | BNC-FL0385 |
| 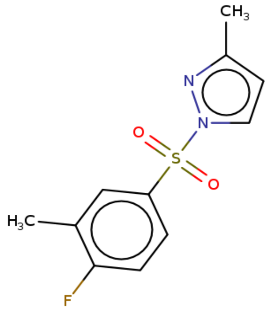 | BNC-FL0506 | 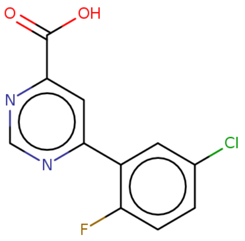 | BNC-FL0209 |
| 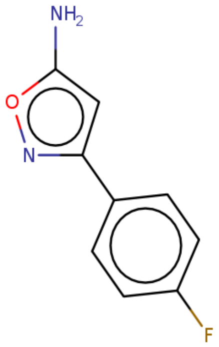 | BNC-FL0770 | 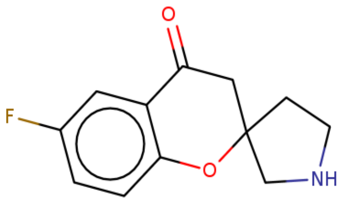 | BNC-FL0555 |

| | | | |
|---|---|---|---|
| 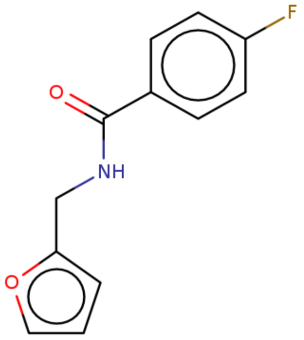 | BNC-FL0470 | 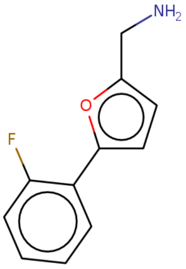 | BNC-FL0772 |
| 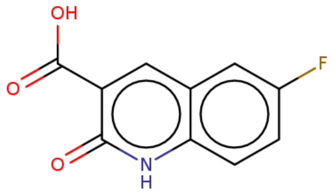 | BNC-FL0219 | 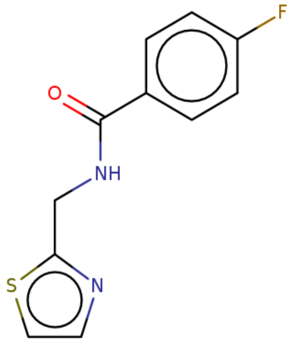 | BNC-FL0501 |
| 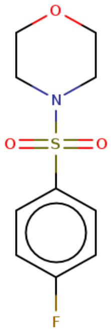 | BNC-FL0315 | 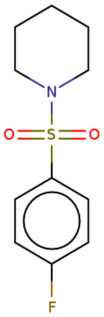 | BNC-FL0363 |

**Table S5** s2m-CoV-2 hits. Fragments that had enhanced $^{19}F$ relaxation in the initial screening.

| | | | |
|---|---|---|---|
| 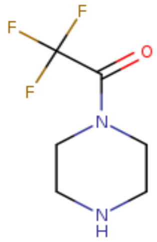 | BNC-TFL103 | 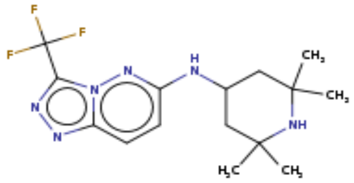 | BNC-TFL384 |
| 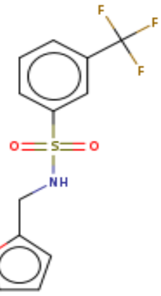 | BNC-TFL192 | 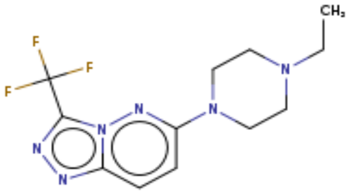 | BNC-TFL389 |
| 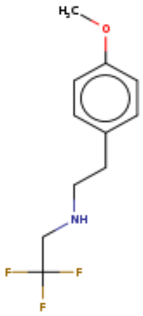 | BNC-TFL326 | 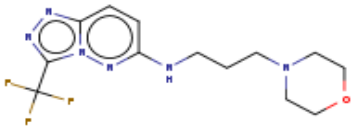 | BNC-TFL371 |
| 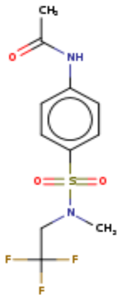 | BNC-TFL352 | 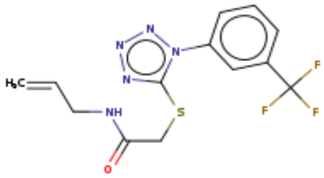 | BNC-TFL405 |

| | | | |
|---|---|---|---|
| 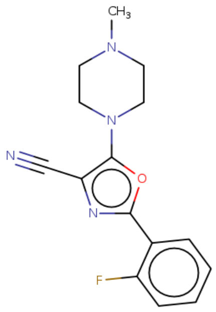 | BNC-FL0757 | 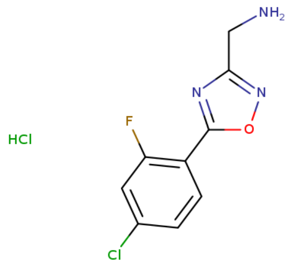 | BNC-FL0051 |
| 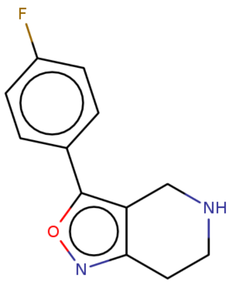 | BNC-FL0123 | 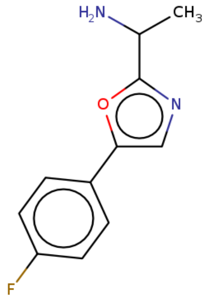 | BNC-FL0092 |
| 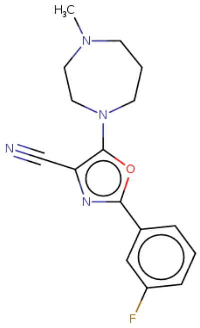 | BNC-FL0726 | 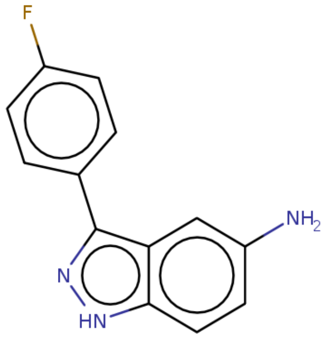 | BNC-FL0096 |
| 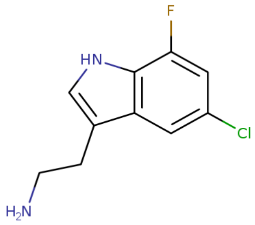 | BNC-FL0049 | 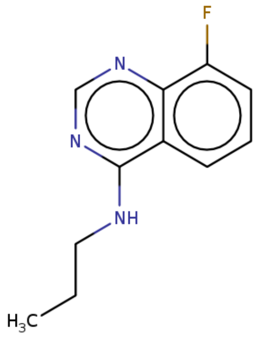 | BNC-FL0496 |

### 2. Supplementary Figures

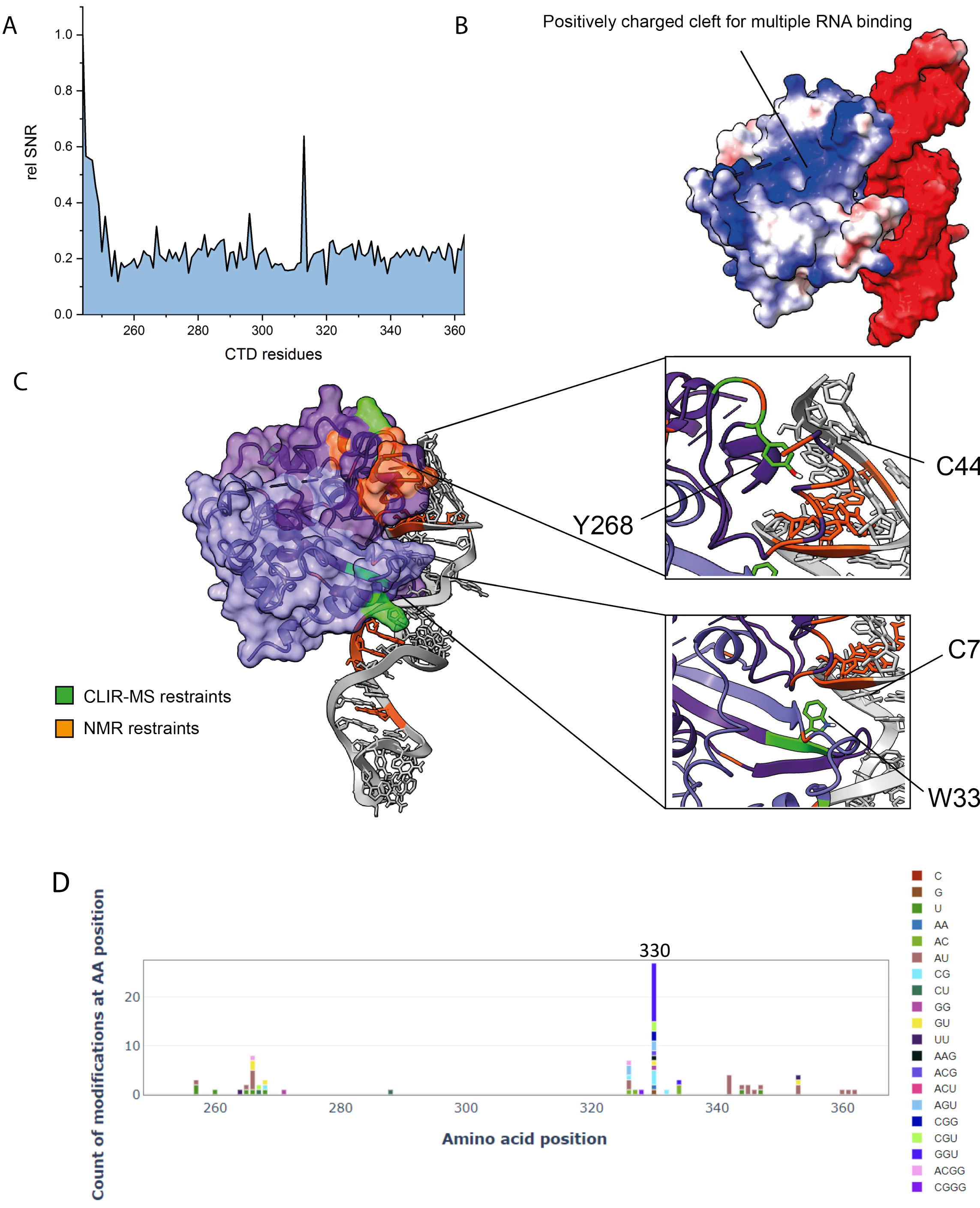

E

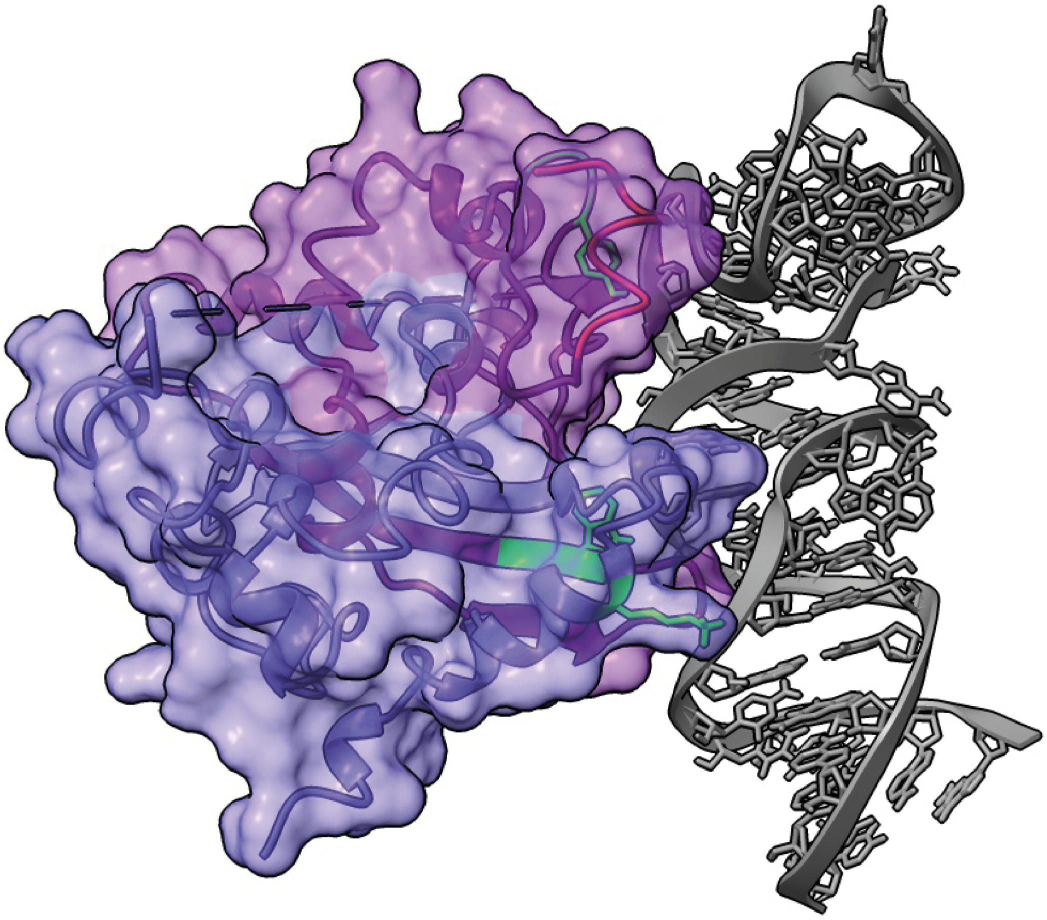

**Figure S1** (A) Signal-to-noise plot of the CTD residues at 0.03 equivalents of s2m-CoV-2. (B) Electrostatic surface of model 1 that highlights a large positively charged cleft still available for RNA binding. (C) Alternative HADDOCK conformation of CTD bound to s2m-CoV-2 calculated using NMR and CLIR-MS restraints. (D) CLIR-MS plot of CTD-s2m-CoV-2 in the presence of Mg. (E) HADDOCK model of the CTD binding the single nucleotide mutant structure obtained by x-ray crystallography (PDB 1XJR).

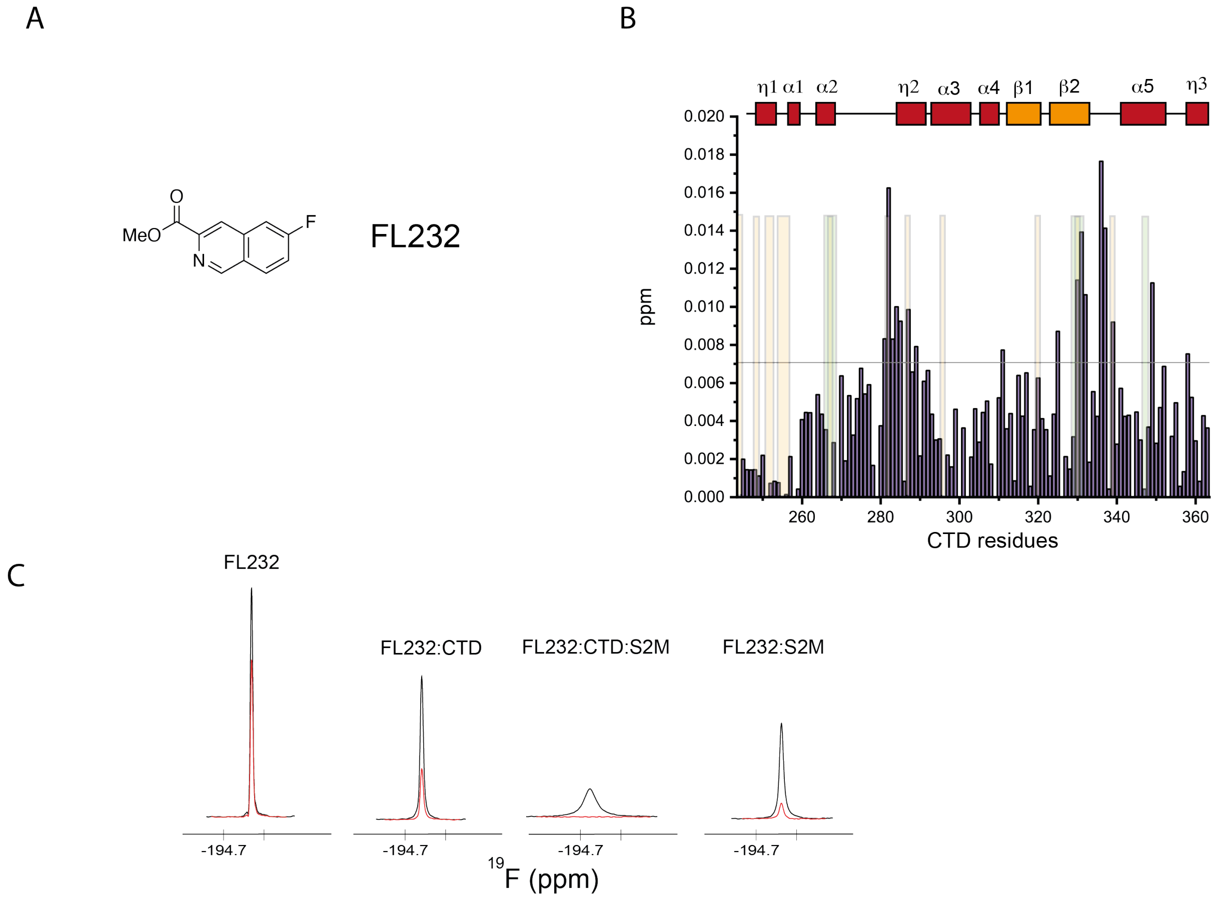


**Figure S2** (A) Structure of FL232. (B) $^{1}$H-$^{15}$N combined chemical shift perturbation plot of the CTD (50 μM) induced by FL232 (700 μM) (orange and green shaded bars function as a reference for perturbed regions of CTD upon s2m-CoV-2 addition detected by NMR and CLIR-MS respectively). (C) $^{19}$F resonances of FL232 in the presence or absence of CTD and s2m-CoV-2. Black: 1D $^{19}$F recorded using 10 ms relaxation delay; Red: 1D $^{19}$F recorded using 200 ms relaxation delay *(*CTD *50 μM,* fragment *750 μM,* RNA*, 20 μM,* 25 mM Na Phos, 50 mM NaCl pH 6, 298 K*)*.

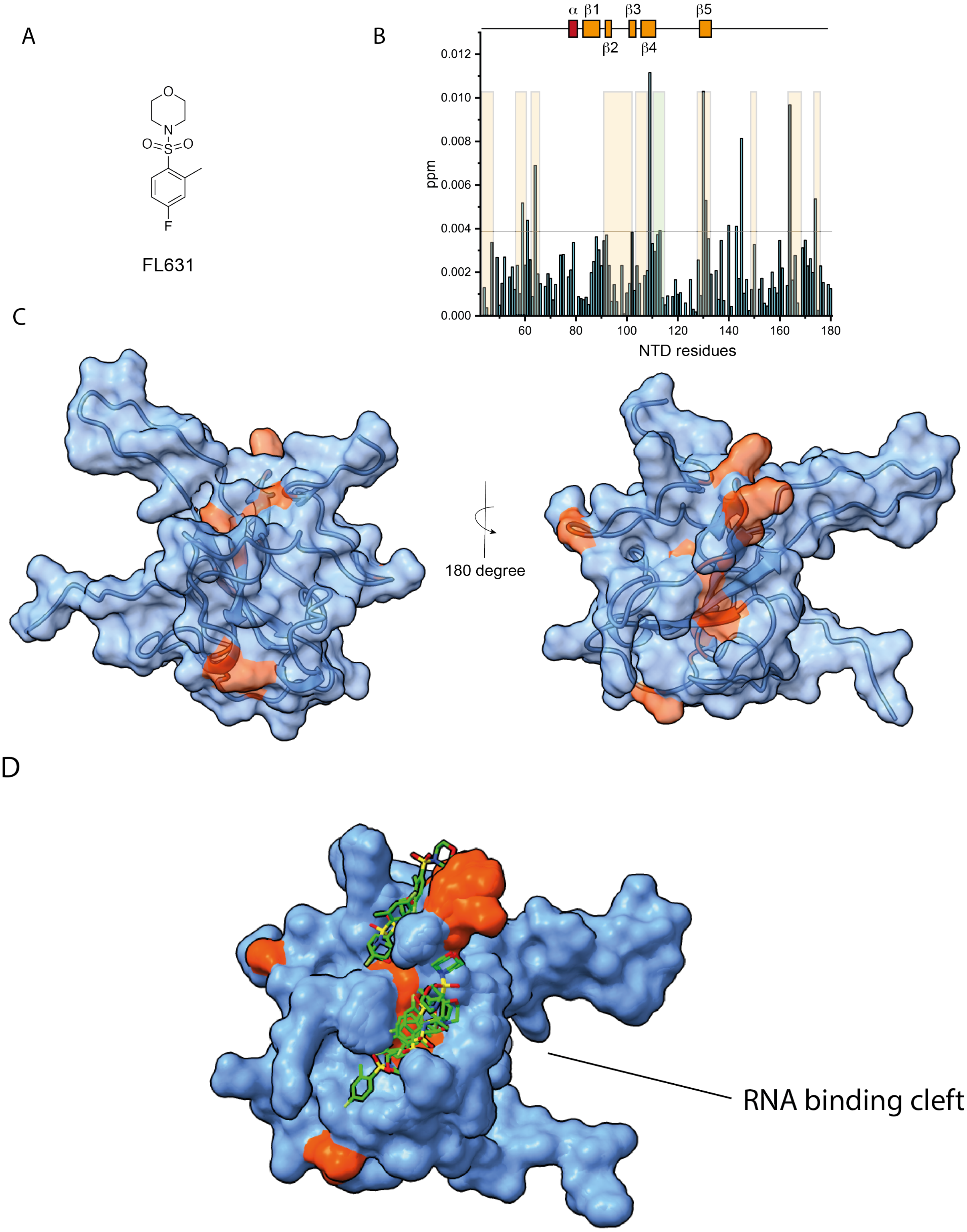


**Figure S3** (A) Structure of a representative sulfonamide fragment FL631. (B) Combined $^{1}H$-$^{15}N$ chemical shift perturbation plot underscoring NTD (50 μM) residues that are shifted upon addition of 15 equivalents of FL631 (750 μM) (50 mM Na Phos, 100 mM NaCl, pH 6.8, 298 K). (C) 3D structure of NTD with highlighted residues (orange) with chemical shift perturbations larger than 2 standard deviations, which suggest that the binding location of FL631 is not at the RNA binding interface. (D) Docking poses obtained from experimentally restrained docking FL631 to NTD.

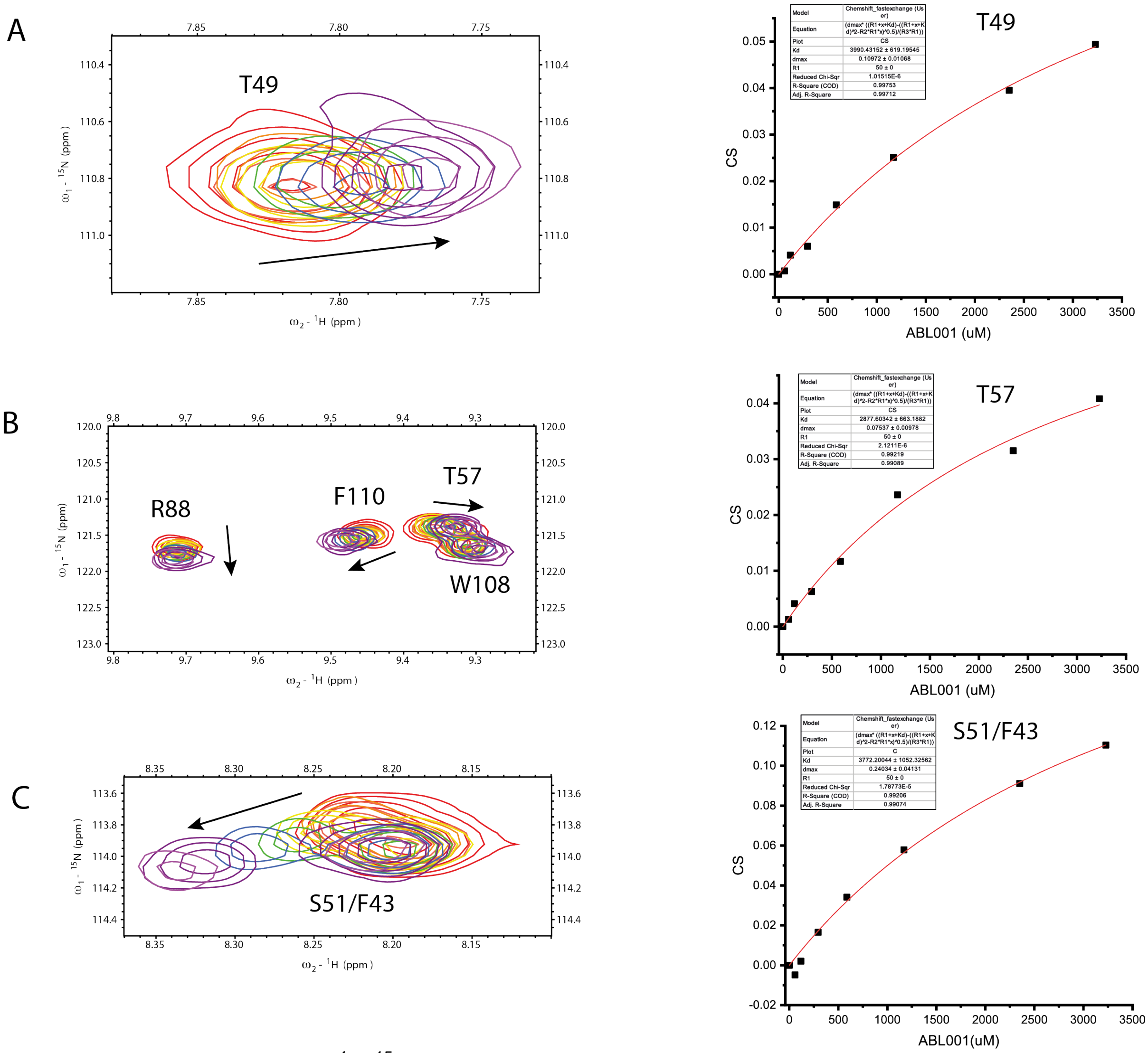


**Figure S4** Zoom of overlayed $^{1}H$-$^{15}N$ HSQC spectra focused on selected peaks showing the titration of ABL001 to NTD. $^{1}H$ perturbation of residues (A) T49, (B) T57, (C) S51/F43 were plotted to estimate the $K_D$ value.

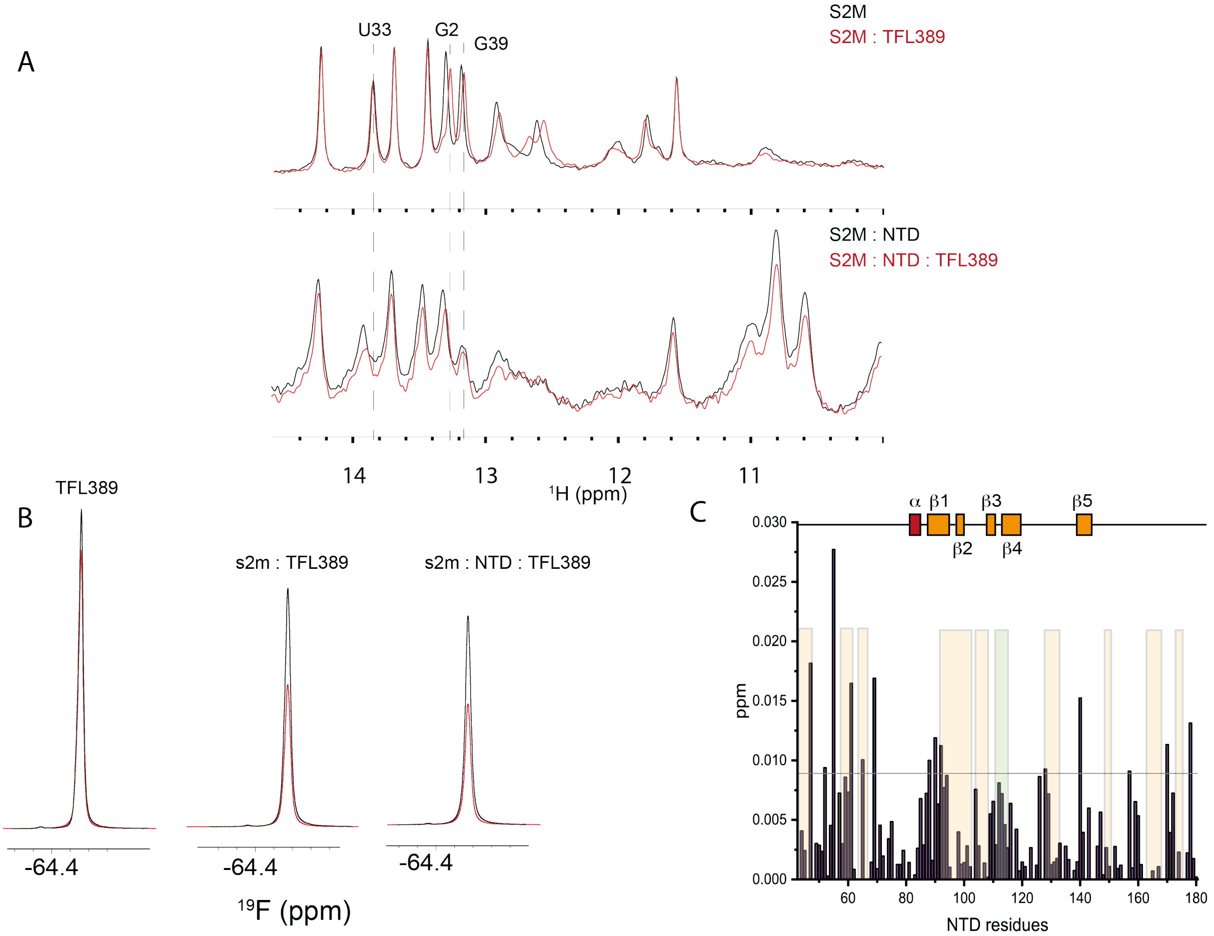


**Figure S5** 1D $^1$H imino of s2m-CoV-2 in the presence or absence of NTD and/or TFL389 in buffer A at 298 K (50 μM s2m-CoV-2, 150 μM NTD, 1.5 mM TFL389). (B) $^{19}$F relaxation of TFL389 under condition B at 310 K in the presence and absence of s2m-CoV-2 and/or NTD. (C) Combined $^1$H-$^{15}$N chemical shift perturbation plot underscoring NTD residues of the NTD-s2m-CoV-2 complex that are shifted upon addition of 15 equivalents of TFL389 (50 mM Na Phos, 100 mM NaCl, pH 6.8, 310 K).

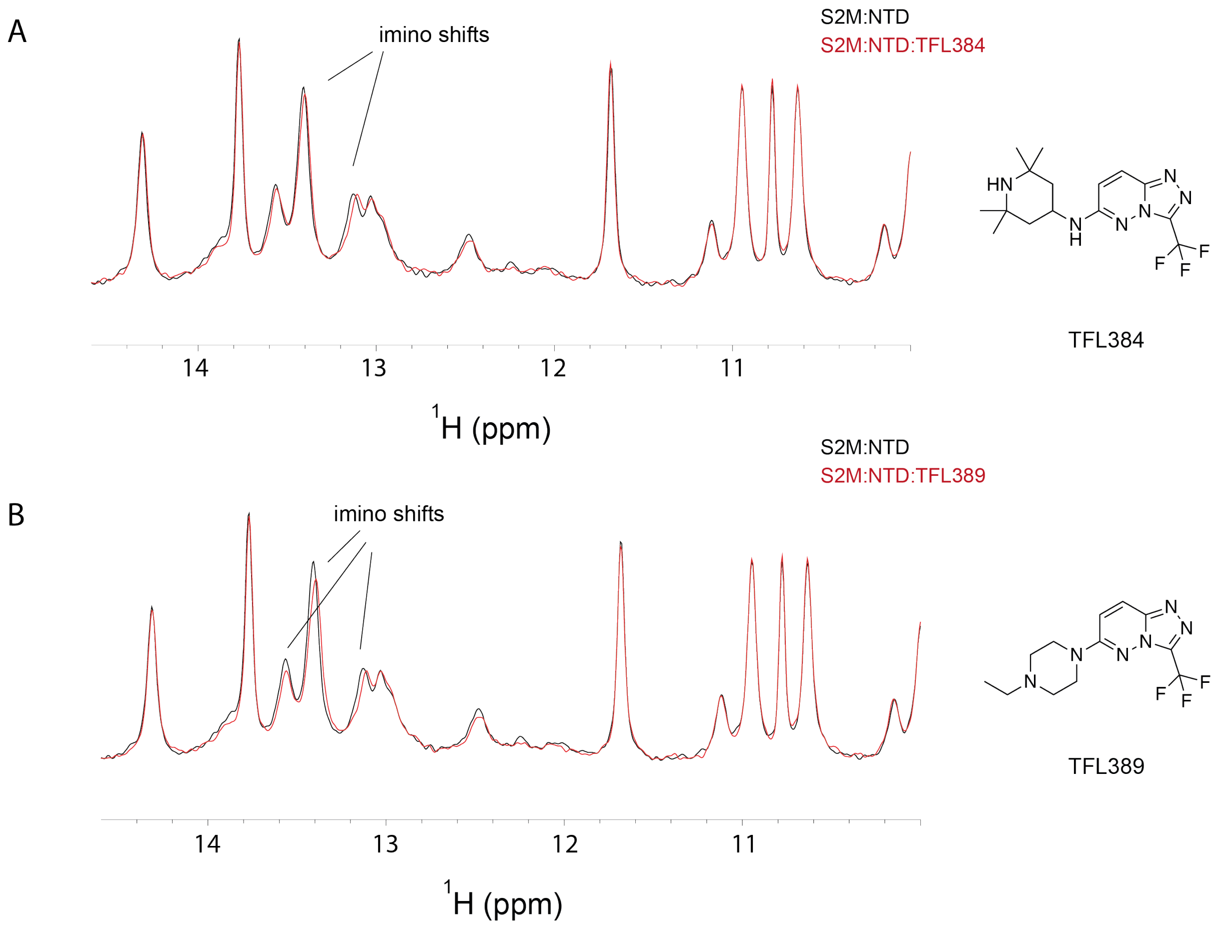


**Figure S6** 1D $^{1}$H SOFAST focused on iminos of s2m-CoV-2-NTD complex (100 μM NTD, 100 μM s2m-CoV-2) in the presence or absence of TFL384 and TFL389 (1.5 mM) in buffer B at 310 K.

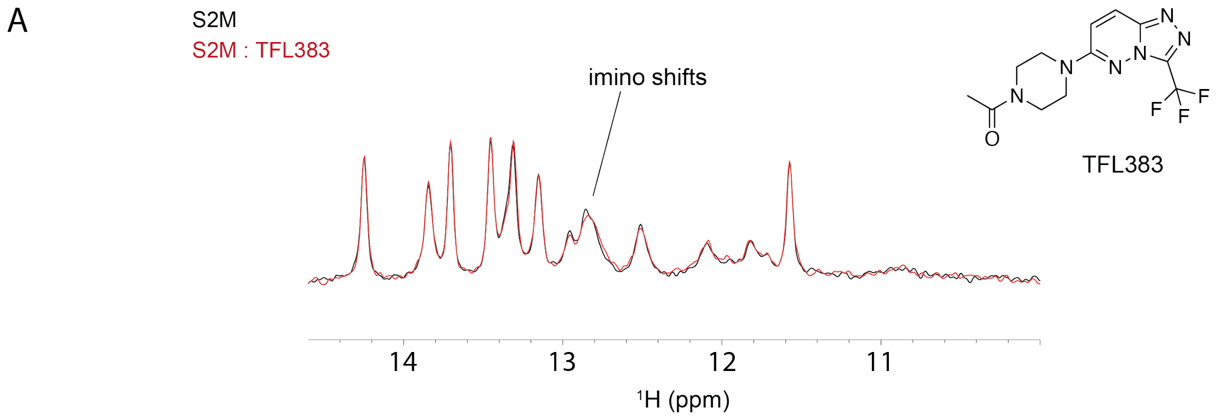


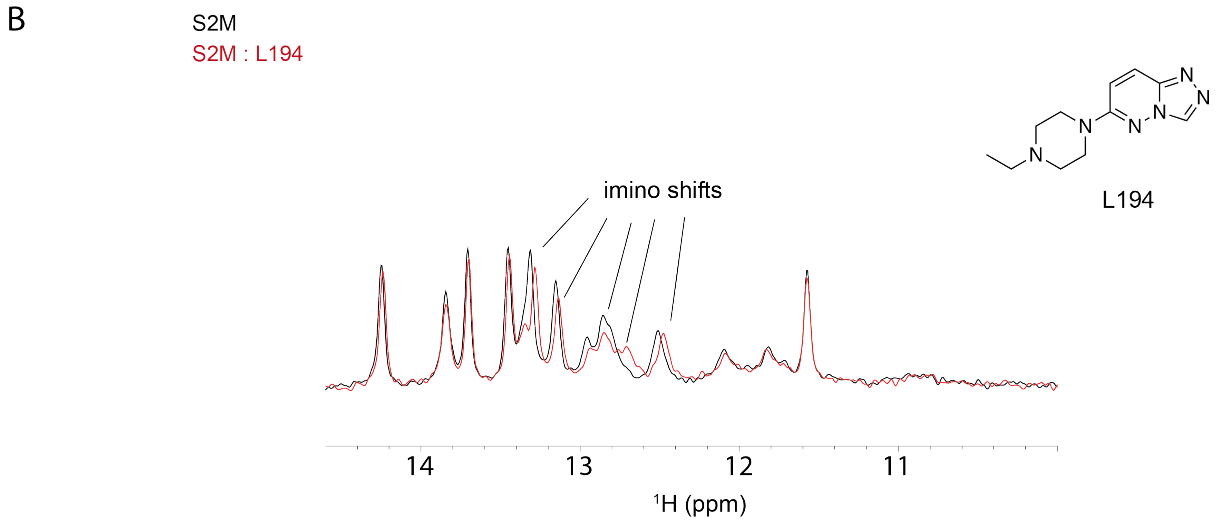


**Figure S7** 1D $^{1}$H SOFAST focused on imino of s2m-CoV-2 (50 μM) in the presence or absence of analogues TFL383 and L194 (1.5 mM) in buffer A at 298 K.

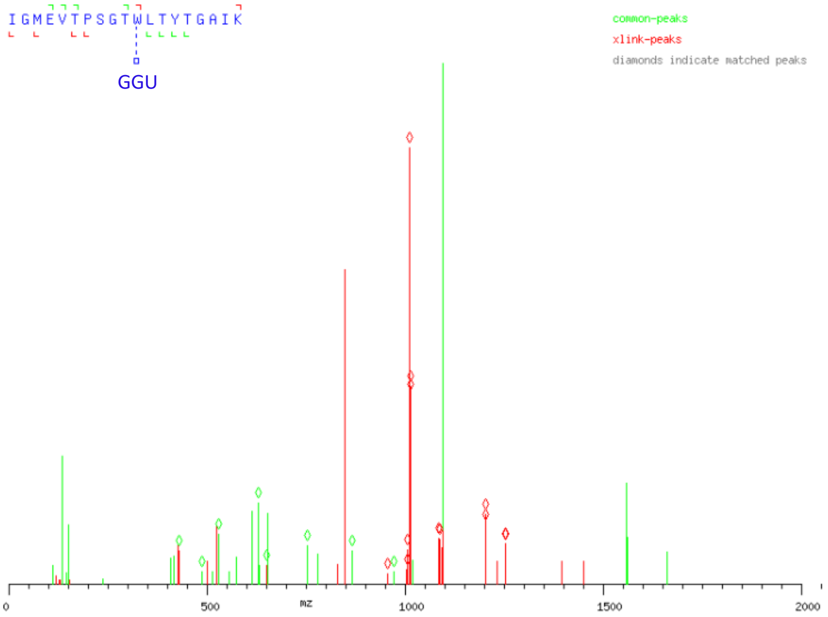


### alpha-chain

| | | | | | | | | | | | | | | | | | | | |
|---|---|---|---|---|---|---|---|---|---|---|---|---|---|---|---|---|---|---|---|
| alpha_common_b_standard_plus1 | 114.09 | 171.11 | 302.15 | 431.20 | 530.26 | 631.31 | 728.37 | 815.40 | 872.42 | 973.47 | - | - | - | - | - | - | - | - | - |
| alpha_common_b_standard_plus2 | 57.55 | 86.06 | 151.58 | 216.10 | 265.64 | 316.16 | 364.69 | 408.20 | 436.71 | 487.24 | - | - | - | - | - | - | - | - | - |
| alpha_common_b_standard_plus3 | 38.70 | 57.71 | 101.39 | 144.40 | 177.43 | 211.11 | 243.46 | 272.47 | 291.48 | 325.16 | - | - | - | - | - | - | - | - | - |
| alpha_xlink_b_standard_plus1 | - | - | - | - | - | - | - | - | - | - | 2171.66 | 2284.74 | 2385.79 | 2548.86 | 2649.90 | 2706.93 | 2777.96 | 2891.05 | 3019.14 |
| alpha_xlink_b_standard_plus2 | - | - | - | - | - | - | - | - | - | - | 1086.33 | 1142.88 | 1193.40 | 1274.93 | 1325.46 | 1353.97 | 1389.49 | 1446.03 | 1510.07 |
| alpha_xlink_b_standard_plus3 | - | - | - | - | - | - | - | - | - | - | 724.56 | 762.25 | 795.94 | 850.29 | 883.97 | 902.98 | 926.66 | 964.35 | 1007.05 |
| AA | I | G | M | E | V | T | P | S | G | T | W | L | T | Y | T | G | A | I | K |
| alpha_common_y_standard_plus1 | - | - | - | - | - | - | - | - | - | - | - | 866.50 | 753.41 | 652.37 | 489.30 | 388.26 | 331.23 | 260.20 | 147.11 |
| alpha_common_y_standard_plus2 | - | - | - | - | - | - | - | - | - | - | - | 433.75 | 377.21 | 326.69 | 245.16 | 194.63 | 166.12 | 130.60 | 74.06 |
| alpha_common_y_standard_plus3 | - | - | - | - | - | - | - | - | - | - | - | 289.50 | 251.81 | 218.13 | 163.77 | 130.09 | 111.08 | 87.40 | 49.71 |
| alpha_xlink_y_standard_plus1 | 3037.15 | 2924.07 | 2867.05 | 2736.01 | 2606.96 | 2507.89 | 2406.85 | 2309.79 | 2222.76 | 2165.74 | 2064.69 | - | - | - | - | - | - | - | - |
| alpha_xlink_y_standard_plus2 | 1519.08 | 1462.54 | 1434.03 | 1368.51 | 1303.99 | 1254.45 | 1203.93 | 1155.40 | 1111.89 | 1083.37 | 1032.85 | - | - | - | - | - | - | - | - |
| alpha_xlink_y_standard_plus3 | 1013.06 | 975.36 | 956.35 | 912.67 | 869.66 | 836.64 | 802.95 | 770.60 | 741.59 | 722.59 | 688.90 | - | - | - | - | - | - | - | - |

**Figure S8** Annotated MS/MS spectrum of the peptide IGMEVTPSGTWLTYTGAIK cross-linked to a GGU trinucleotide indicating the cross-linking to the W330 amino acid position.

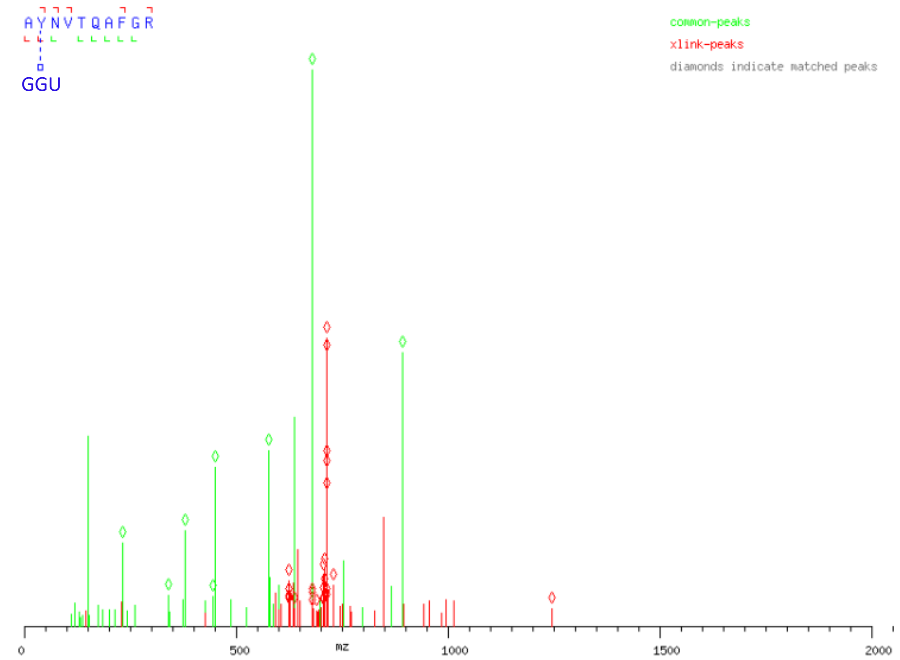


### alpha-chain

| | | | | | | | | | | |
|---|---|---|---|---|---|---|---|---|---|---|
| alpha_common_b_standard_plus1 | 72.04 | - | - | - | - | - | - | - | - | - |
| alpha_common_b_standard_plus2 | 36.53 | - | - | - | - | - | - | - | - | - |
| alpha_common_b_standard_plus3 | 24.69 | - | - | - | - | - | - | - | - | - |
| alpha_xlink_b_standard_plus1 | - | 1247.22 | 1361.27 | 1460.33 | 1561.38 | 1689.44 | 1760.48 | 1907.55 | 1964.57 | 2120.67 |
| alpha_xlink_b_standard_plus2 | - | 624.12 | 681.14 | 730.67 | 781.20 | 845.22 | 880.74 | 954.28 | 982.79 | 1060.84 |
| alpha_xlink_b_standard_plus3 | - | 416.41 | 454.43 | 487.45 | 521.13 | 563.82 | 587.50 | 636.52 | 655.53 | 707.56 |
| AA | A | Y | N | V | T | Q | A | F | G | R |
| alpha_common_y_standard_plus1 | - | - | 892.46 | 778.42 | 679.35 | 578.31 | 450.25 | 379.21 | 232.14 | 175.12 |
| alpha_common_y_standard_plus2 | - | - | 446.74 | 389.71 | 340.18 | 289.66 | 225.63 | 190.11 | 116.57 | 88.06 |
| alpha_common_y_standard_plus3 | - | - | 298.16 | 260.15 | 227.12 | 193.44 | 150.75 | 127.08 | 78.05 | 59.05 |
| alpha_xlink_y_standard_plus1 | 2138.68 | 2067.64 | - | - | - | - | - | - | - | - |
| alpha_xlink_y_standard_plus2 | 1069.84 | 1034.33 | - | - | - | - | - | - | - | - |
| alpha_xlink_y_standard_plus3 | 713.57 | 689.89 | - | - | - | - | - | - | - | - |

**Figure S9** Annotated MS/MS spectrum of the peptide AYNVTQAFGR cross-linked to a GGU trinucleotide indicating the cross-linking to the Y268 amino acid position.
